# Loss of NMDA receptor function during development results in decreased KCC2 expression and increased neurons in the zebrafish forebrain

**DOI:** 10.1101/2023.08.25.554812

**Authors:** Amalia J. Napoli, Stephanie Laderwager, Josiah D. Zoodsma, Bismi Biju, Olgerta Mucollari, Sarah K. Schubel, Christieann Aprea, Aaliya Sayed, Kiele Morgan, Annelysia Napoli, Stephanie Flanagan, Lonnie P. Wollmuth, Howard I. Sirotkin

**Author notes:** Co-senior authors (Lonnie P. Wollmuth, Howard I. Sirotkin). **Address for correspondence:**Dr. Howard I. Sirotkin Depts. of Neurobiology & Behavior Stony Brook University Stony Brook, New York 11794-5230.

## Abstract

Developmental neurogenesis is a tightly regulated spatiotemporal process with its dysregulation implicated in neurodevelopmental disorders. NMDA receptors are glutamate-gated ion channels that are widely expressed in the early nervous system, yet their contribution to neurogenesis is poorly understood. Notably, a variety of mutations in genes encoding NMDA receptor subunits are associated with neurodevelopmental disorders. To rigorously define the role of NMDA receptors in developmental neurogenesis, we used a mutant zebrafish line (*grin1*^-/-^) that lacks all NMDA receptors yet survives to 10 days post-fertilization, offering the opportunity to study post-embryonic neurodevelopment in the absence of NMDA receptors. Focusing on the forebrain, we find that these fish have a progressive supernumerary neuron phenotype confined to the telencephalon at the end of embryonic neurogenesis, but which extends to all forebrain regions during postembryonic neurogenesis. This enhanced neuron population does not arise directly from increased numbers or mitotic activity of radial glia cells, the principal neural stem cells. Rather, it stems from a lack of timely maturation of transit-amplifying neuroblasts into post-mitotic neurons, as indicated by a decrease in expression of the ontogenetically-expressed chloride transporter, KCC2. Pharmacological blockade with MK-801 recapitulates the *grin1*^-/-^ supernumerary neuron phenotype, indicating a requirement for ionotropic signaling. Thus, NMDA receptors are required for suppression of indirect, transit amplifying cell-driven neurogenesis by promoting maturational termination of mitosis. Loss of suppression results in neuronal overpopulation that can fundamentally change brain circuitry and may be a key factor in pathogenesis of neurodevelopmental disorders caused by NMDA receptor dysfunction.

## INTRODUCTION

Precise spatiotemporal regulation of developmental neurogenesis is required for proper brain cytoarchitecture. Neurons are generated directly from multipotent neural stem cells (NSC) and indirectly from their unipotent progeny, the neural progenitor cells (NPC) [1–3]. NPCs are neuronally fate-restricted. Before maturing into postmitotic neurons, they function as intermediate progenitor cells and other transit amplifying cell populations (tAC) [4–6]. Neural stem and progenitor cells (NSPC) generate neurons in different brain regions at different times throughout embryological and postnatal development [7]. Fate specification and density of these neurons is largely dictated by NSPC source, time, and location of their birth [8,9]. Disruption of this regulatory balance in neurogenesis is associated with neurodevelopmental disorders (NDD) including autism spectrum disorder (ASD) [10,11].

NMDA receptors are glutamate-gated ion channels that are widely expressed throughout the brain. They play essential ionotropic and non-ionotropic signaling roles at excitatory synapses in the mature brain [12], and during neurodevelopment in cells that do not have synapses [13]. NMDA receptors are heterotetramers that contain two obligate GluN1 subunits, encoded by the *GRIN1* gene and, typically, some combination of two GluN2A-2D subunits encoded by *GRIN2A-2D*. Notably, mutations in *GRIN* genes are associated with NDD including ASD, epilepsy, and schizophrenia [14–16] though their mechanisms of pathogenesis are unknown. Many of the neuronal subtypes implicated in NDDs are generated or mature postnatally [17,18]. Thus, understanding the regulatory role of NMDA receptors during all stages of developmental neurogenesis will shed light on NDD etiology.

NMDA receptors are expressed in all NSPC [19,20]. NSC-driven direct neurogenesis and tAC-driven indirect neurogenesis are spatially and temporally balanced to appropriately build different brain regions [21]. The suppression of indirect neurogenesis is tied to maturation of tACs to a post mitotic state with upregulation of the neuron-specific potassium chloride cotransporter (KCC2) as one of the earliest mileposts of this maturation [22]. The specific role of NMDA receptors in regulating direct and indirect neurogenesis is unclear. Further highlighting the complex role of NMDA receptors in neurogenesis is that they have been proposed to both suppress [23–26] and promote [27–31] neurogenesis. These differences are presumably due to variations in specific NSPC focus and stages of neurogenesis investigated, each of which differentially contribute to direct and indirect neurogenesis.

*Grin1* knockout in rodents is perinatal lethal [32]. NMDA receptors are required for the perinatal suckling response [33], which may contribute to the lethality of *Grin1* knockouts. On the other hand, zebrafish with both paralogous *grin1* genes disrupted (*grin1*^-/-^ fish) survive to 10 days post fertilization (dpf) [34], presumably because suckling is not required. The molecular and cellular mechanisms of neurodevelopment are highly conserved across vertebrates making zebrafish a powerful experimental model for humans. In addition, neurodevelopment in zebrafish is very rapid; 3 dpf marks the beginning of postembryonic neurogenesis, therefore, *grin1*^-/-^ fish survive far beyond the comparable survival stage in rodents. Since GluN1 is obligatory, *grin1*^-/-^ fish lack all NMDA receptors. Thus, this *in vivo* model provides a unique window into postembryonic neurogenesis in the context of complete NMDA receptor loss of function.

Here, we focused our experiments on the forebrain because it primarily undergoes programmed neurodevelopment, as opposed to other brain regions, such as the midbrain optic tectum, where neurodevelopment is strongly activity-dependent [35,36]. We find that loss of NMDA receptors results in supernumerary forebrain neurons in zebrafish. This phenotype is first evident in the telencephalon at the end of embryonic development but progresses both in magnitude and spatially to encompass the entire forebrain during postembryonic neurogenesis. We determined this dysregulation is not due to changes in direct neurogenesis but rather delayed maturation of tACs, as evidenced by reduced expression of KCC2. Maturational delay prolongs tAC proliferation and inappropriately amplifies the neuronal pool. Thus, NMDA receptors have a key role in suppressing indirect neurogenesis by promoting maturation of tAC neuroblasts into post-mitotic neurons. These findings provide a framework in which to study the effects of loss- and gain-of-function mutations in NMDA receptor subunits and their effects on neurogenesis, KCC2 expression, and NDD pathogenesis.

## MATERIALS AND METHODS

### Zebrafish Husbandry

Zebrafish were housed in a dedicated facility at 28.5⁰ C and were maintained under a 13/11 light/dark cycle and a diet of artemia and GEMMA micropellets. All experimental work contained herein was approved and conducted in accordance with the Stony Brook University Institutional Animal Care and Use Committee (IACUC).

### Specimen Generation

All embryos were generated from natural crosses. For all experiments, unless otherwise indicated, *grin1*^-/-^ fish were generated from parents harboring heterozygous *grin1a* mutations and homozygous *grin1b* mutations [34]. After fixation, approximately 1 mm of the larval tail was removed with a microdissection scissor for genotyping. Samples were boiled in lysis buffer, digested with Proteinase K (BioVision) and genotyped by PCR using the following primers: *grin1a*: forward (TGGGCTGGCTTCGCTATGAT) and reverse (GGGTCGTTGATGCCGGTGAT); *grin1b*: forward (GGTGCCCCTCGGAGCTTTTC) and reverse (GGAAGGCTGCCAAATTGGCAGT). Controls for each assay were wild type (hybrid wild-type strain consisting of Tubingen long-fin and Brian’s wild type (TLB)) fish time-matched for each clutch of mutant fish. At the stages of development investigated, sex is indeterminant in zebrafish and was, thus, not considered as a variable.

### Histology

Zebrafish larvae were fixed in neutral buffered formalin overnight at room temperature (RT). Larvae were then rinsed three times in 70% ethanol and left in the final rinse overnight at RT. Larvae were mounted in 1% High EEO agarose (Crystalgen) molds [37]. Embedded fish were dehydrated in progressive alcohols and cleared in xylene under 15 inHg vacuum pressure at RT. Embedded fish were infused with paraffin (Leica Microsystems FormulaR) under 15 inHg at 60⁰ C. Paraffin blocks were sectioned coronally at 5 µm on a Spencer rotary microtome. Slide mounted sections were warmed overnight, cleared in xylene, progressively rehydrated to tap water, stained in cresyl violet (Sigma-Aldrich), and progressively dehydrated to xylene. Slides were cover glass mounted using DPX mounting medium (Sigma-Aldrich). Slides were imaged on an Olympus VS120 slide scanning microscope.

### Immunohistochemistry (For antibody, Research Resource Identifiers, and protocol specifics see Table 1)

#### For immunohistochemistry (IHC) performed on cryotomy sections

Zebrafish larvae were fixed in 4% PFA for 2-3 hours at RT, rinsed three times in PBS and incubated in 30% sucrose water overnight at 4⁰ C, then mounted in 1% High EEO agarose molds made with 30% sucrose (molds previously described in histology methods). Agarose molds were embedded in Tissue-Tek O.C.T compound (Sakura), frozen, and coronally sectioned at a thickness of 10 µm on a Leica CM1950 cryostat. Slides with mounted tissue were warmed for 15 minutes for tissue adherence, triple rinsed (5 minutes each rinse) in PBS, incubated in antigen retrieval solution, then triple rinsed in PBS, followed by two rinses in PBS with 0.3% TritonX (ThermoScientific)(PBSTx) before blocking and primary antibody.

**Table 1.** List of abbreviations.

| *Abbreviation* | *Term* |
| --- | --- |
| ac | Anterior Commissure |
| ED | Eye diameter |
| EE | Distance from eye to ear |
| EmT | Thalamic eminence |
| GFAP | Glial Fibrillary Acidic Protein |
| Ha | Habenula |
| Hr | Rostral Hypothalamus |
| iPC | Intermediate Progenitor Cell |
| KCC2 | Potassium-chloride transporter member 5 |
| NEC | Neuroepithelial Cell |
| NPC | Neural Progenitor Cell |
| NSC | Neural Stem Cell |
| NSPC | Neural Stem and Progenitor Cell |
| Pa | Anterior Pallium |
| PCNA | Proliferating Cell Nuclear Antigen |
| Pm | Mid Pallium |
| Po | Preoptic area |
| PSA-NCAM | Polysialylated Neural Cell Adhesion Molecule |
| PT | Posterior Tuberculum |
| PVZ | Periventricular Zone |
| RGC | Radial Glia Cell |
| Sd | Dorsal Aspect of the Subpallium |
| T | Telencephalon |
| tAC | Transit Amplifying Cell |
| TeO | Optic Tectum |

**Table 2.** List of Antibodies.

| Antibody | Manufacturer | Host | Concentration | Antigen Retrieval/<br>Pretreatment | Blocking Solution | Antibody<br>Incubation<br>Solution |
| --- | --- | --- | --- | --- | --- | --- |
| anti-GFAP<br>(zrf1) | ZIRC<br>RRID:<br>AB_10013806 | M | 1:75 | SCB pH 8.5 at 83° C<br>for 30 minutes | 6% donkey serum in<br>0.3% PBSTx | Same as block |
| anti-<br>cleaved<br>Caspase-3 | Cell Signaling<br>RRID:<br>AB_2341188 | R | 1:500 | Iced 100% Methanol<br>PDT | 10% heat-inactivated<br>fetal bovine serum + 2%<br>bovine serum albumin in<br>PBST | Same as block |
| anti-PCNA | GeneTex<br>RRID:<br>AB_11161916 | R | 1:500 | SCB pH 8.5 at 83° C<br>for 30 minutes | 6% donkey serum in<br>0.3% PBSTx | Same as block |
| anti-PCNA | Sigma Aldrich<br>RRID:<br>AB_477413 | M | 1:250 | SCB pH 8.5 at 80° C<br>for 30 minutes | 6% donkey serum in<br>0.1% bovine serum<br>albumin solution made<br>with 0.3% PBSTx | Same as block |
| anti-PSA-NCAM | Millipore<br>RRID:<br>AB_95211 | M | 1:500 | 3% hydrogen peroxide<br>in 100% methanol | 1% bovine serum<br>albumin in 0.3% PBSTx | Same as block |
| anti-SLC12A5<br>(KCC2) | Sigma Aldrich<br>RRID:<br>AB_2188801 | R | 1:250 | SCB pH 8.5 at 80° C<br>for 30 minutes | 6% donkey serum in<br>0.1% bovine serum<br>albumin solution made<br>with 0.3% PBSTx | Same as block |
| anti-GluN1 | Synaptic<br>Systems<br>RRID:<br>AB_887750 | M | 1:200 | 1N HCL at 37° C for<br>20 minutes<br>Normalize in Borate<br>Buffer pH 8.5 | 6% donkey serum in<br>0.3% PBSTx | 0.1% BSA in<br>PBS |

Antibodies and immunohistochemical procedures used for each assay.

Tissue was blocked in animal serum (donkey, Millipore; bovine serum albumin, Lampire) in PBSTx for 1 hour at RT, then incubated in primary antibody at 4⁰ C for 24-36 hours (see Table 1). Tissue was triple rinsed in PBS followed by two rinses in PBSTx, then incubated in secondary antibody for 3-4 hours at RT. All secondary antibodies used were Alexa Fluor 488 or 594, donkey anti-mouse or rabbit (Invitrogen) as required for specific primary antibody. Slides were cover glass mounted using DAPI Fluoromount (Southern BioTech). Images were acquired on an Olympus VS120 slide scanning microscope and an Olympus FV1000. In all experiments, larvae used for quantitative comparison received the same preparation and treatment and the same imaging parameters.

#### For IHC performed on whole-mount zebrafish larvae

Zebrafish larvae were fixed in 4% PFA for 2-3 hours at RT, rinsed 2 times in PBS and once in 0.1% TWEEN (Sigma-Aldrich) in PBS (PBST), then depigmented in a solution of 50 mM potassium hydroxide in 1.5% hydrogen peroxide for 30 minutes at RT, then washed in PBS. Larvae were then incubated in ice-cold 100% methanol for 2 hours at -20⁰ C to permeabilize, followed by incubation in a mixture of PBST +0.3% Triton-X and 1% DMSO (PDT) applied twice for 30 minutes at RT. Larvae were blocked for 1 hour at RT, then incubated in primary antibody overnight at 4⁰ C (see Table of Antibodies). Larvae were then blocked again for 1 hour at RT and then incubated in secondary antibody for 2 hours at RT. Embryos were deyolked and mounted dorsally on slides with Krystalon mounting medium (Sigma-Aldrich). Images were acquired on an Olympus VS120 slide scanning microscope.

### Quantification and Analysis

#### For all analytical methods described below

Images for analysis were selected by anatomical levels, which were landmarked and standardized anatomically after Mueller & Wullimann (2016) [38]. Where relevant, anatomical levels are numbered for each postembryonic stage according to Mueller & Wullimann, Chapter 2: Hu Protein Expression (2016) [38]. All analyses were performed by investigators blinded to genotype to prevent bias.

### Morphometric Analysis

#### For direct morphometric measurements

Measurements were performed on 5 dpf larval zebrafish brain sections prepared according to histological methods above. Three levels of the forebrain were assayed for dorsoventral height, mediolateral width, and hemispheric area; the telencephalon of the anterior forebrain was assessed at the level of the anterior commissure, the mid forebrain was assessed at the posterior telencephalon/diencephalic boundary, and the posterior forebrain was assessed in the diencephalon, at the level of the rostral hypothalamus anterior to the optic tectum. Dorsolateral measurements were made at the midline of the brain. Mediolateral measurements were made at the widest aspect of the brain for each anatomical level. Hemispheric area was assessed using the dorsoventral midline as the inner boundary and tracing along the parenchymal-meningeal boundary of one hemisphere.

#### For external proxy measurements

Measurements for brain size, as described and validated by Naslund (2014) [39], were performed on live larvae at 6 dpf. Larvae were anesthetized with a 0.4% stock solution of Tricaine (Sigma-Aldrich) added dropwise to egg water media (0.3 g/L Instant Ocean Sea Salts) in a 50 mm petri dish then mounted in 3% methylcellulose. Dorsal and lateral images were acquired on a Zeiss SteREO Discovery microscope then processed in Fiji (ImageJ) software [40]. Linear measurements were made of external landmarks that correspond to internal brain size as follows: eye diameter (ED) was measured in lateral view from dorsal to ventral outer margins of the retinal epithelium; distance from eye to ear (EE) was measured in lateral view from the dorsal most border of the retinal epithelium to the anterior most border of the otic vesicle; size of optic tectum (TeO) was measured in dorsal view, mediolaterally across the full extent of body tissue directly posterior to and in apposition to the eyes.

### Corrected Fluorescence Intensity (CFI) Analysis

All fluorescence intensity analysis was performed on 10 µm coronal sections and conducted in Fiji [40]. Images were acquired on an Olympus VS120 slide scanning microscope at the same scanning parameters for all specimens within each analysis group. For all types of CFI analysis (total brain, line, and cell) background fluorescence was determined as follows: mean fluorescence intensity measurements were recorded from three sampled regions with no expression for the fluorophore being analyzed. These measurements were obtained from the same section on which the CFI analysis was performed and were selected from within the analyzed region. The three background measurements were averaged to generate a mean fluorescence background value.

#### For GFAP, PCNA, and PSA-NCAM

The forebrain analysis was divided into three gross anatomical regions, the precommissural telencephalon, postcommissural telencephalon, and diencephalon. The precommissural telencephalon sampling was bounded anteriorly at the level of insertion of the olfactory bulb into the pallium and posteriorly at the anterior commissure. The postcommissural telencephalon sampling was bounded anteriorly by the anterior commissure and posteriorly by the caudal-most aspect of the pallium at the boundary with diencephalic territories. Both pre-commissural and post-commissural telencephalon were sampled in the full dorsal to ventral aspect. The diencephalic sampling was bounded anteriorly at the border with the caudal pallium, posteriorly at the rostral hypothalamus, and dorsally at the griseum tectale/fasciculus retroflexus.

#### For each analysis reported in gross anatomical region box plots

At stage 3 dpf, three to five individual atlas levels were assayed within each region; at stage 5 dpf, four to seven individual atlas levels were assayed within each region. The CFI results were averaged for each specimen across the sampled anatomical levels within each of the three gross anatomical regions. Box plots represent the minimum, first quartile, median, third quartile, and maximum values of the data. Scatter represents the average value for each fish, of each genotype, within each gross anatomical region.

#### For each analysis reported in line graphs

Points represent the average for each genotype at each anatomical level assayed. *t-test* was performed as a comparison of the aggregate mean CFI across the forebrain for each group.

#### For corrected total brain fluorescence intensity analysis

Measurements were acquired as follows. The brain was circumscribed; total brain area and integrated fluorescent density across that area was measured. Mean background fluorescence was calculated as described above. The corrected fluorescent intensity of the brain was then calculated according to the following formula:

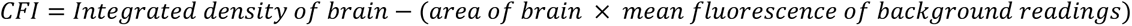

#### For KCC2 line analysis

Fluorescence was measured in each pixel across a transect line to provide a spatial resolution to expression (line analysis). Line analysis was performed in the dorsal aspect of the anterior telencephalon through the pallium and in the diencephalon dorsally through the dorsal thalamus, and ventrally through the posterior tuberculum. To standardize all measurements in all specimens to the individual’s anatomy, the transect was drawn from the midline proliferative zone four cell lengths down from the griseum tectale (dorsal transect) and four cell lengths down from the boundary between the dorsal thalamus and the posterior tuberculum (ventral transect). Transects extended laterally into the parenchyma to a parasagittal plane at the level of the apex of the fasciculus retroflexus just superior to the pretectum. Mean fluorescence for each pixel across the line was measured and corrected for background by subtracting the average background fluorescence (see above). To adjust for intraspecific variation between specimen brains and allow for accurate comparison of distance to initiation of expression, the pixel length was expressed as a percentage of transect. Averages were created in Igor Pro (WaveMetrics) by binning line percentages into 0.5% bins and averaging across each bin for each transect and genotype.

#### For corrected cell fluorescence

For each specimen and proximal to the transects described above, sets of three cells were sampled, one each in an abventricular, parasagittal, and peri-neuropilar location respectively. Locations of cells were standardized for distance from the periventricular zone (PVZ) for each specimen. Cells were circumscribed, mean fluorescence for PCNA and KCC2 was measured, and CFI was determined for each marker by subtracting the average background mean fluorescence (see above) for each fluorophore, respectively.

#### For KCC2 brain CFI

Each specimen was analyzed at the level of the anterior commissure in 10 µm coronal sections. The whole brain at that level was circumscribed, integrated density and background mean fluorescence were measured, and CFI was calculated using the formula above. The CFI of the anterior commissure was similarly assayed and then subtracted from the CFI of the whole brain and normalized to the amount of DAPI^+^ cells to determine the KCC2 expression proportionate to the number of neurons in the cell-rich portion of the tissue.

### Cell Quantification

5 µm coronal paraffin sections were prepared and Nissl stained according to histological methods above. All cell quantification was conducted in Fiji [40]. Distance between sections quantified was 50-60 µm, depending on anatomical distance between cell populations/anatomical regions, to ensure no double counting of cells. Counting was performed by investigators blinded to genotype to prevent bias.

#### For total hemispheric cell counts

Three regions of the forebrain, the anterior, mid, and posterior, were assayed for total hemispheric cell counts. Regions of the forebrain and the brain hemispheres were determined as described above in direct morphometric measurements.

#### For cell density analysis

Cell populations were identified by anatomical means; boxes were drawn in each cell population and area of the box measured. Cells were counted within the box; transected cells were counted on two adjacent sides of the box and excluded on the other two sides. Cell counts were then divided by box area.

#### For PCNA^+^ cell analysis

10 µm coronal sections were analyzed at representative anatomical levels within each of the three gross anatomical regions described previously: precommissural and postcommissural telencephalon, and diencephalon. NSCs and tACs were identified by location (PVZ for NSCs and abventricular for tACs), somal shape (ovoid for RGCs and rounded for tACs) and corroborated with GFAP^+^ status in NSCs.

### Pharmacological Manipulation

Wild-type zebrafish eggs were placed in 30 mm petri dishes filled with 5 mL of egg water (0.3 g/L Instant Ocean Sea Salts) 20 eggs per plate. At 23 hours post fertilization (hpf), embryos were manually dechorionated and placed in either 5 mL of egg water (control group) or 5 mL of 40 µM MK-801(Fisher Scientific) in egg water, which was determined as optimal dosage for effectiveness and viability. Every 24 hours each group’s fluid was replenished.

### Statistical Analysis

All statistical analysis was conducted using R Statistical Software (v4.2.0: R Core Team) [41] with plots generated using ggplot2 (v3. 3.3: Wickham, 2016) [42] and patchwork [43] packages. Equivalence of variance was assessed using an *F*-test, after which statistical significance was determined using either two sample Student’s or Welch’s t-tests assuming equal or unequal variances, respectively, according to the result of the *F*-test for variance. All statistical tests used an *a priori* significance level of 0.05. Outliers were calculated as any data point falling more than 1.5 times the interquartile range above the upper or below the lower quartile. Effect size was calculated and reported using Cohen’s d.

## RESULTS

To determine the role of NMDA receptors (NMDAR) in neurogenesis, we contrasted wild-type with *grin1^-/-^*fish, which lack GluN1 [34]. Because GluN1 is an obligatory subunit, *grin1^-/-^* fish lack all NMDA receptor function. These fish, therefore, provide a unique opportunity to examine the role of NMDA receptors in postembryonic neurodevelopment.

### *grin1*^-/-^ fish do not show gross morphological changes in brain structure

In humans, *GRIN* variants are often associated with gross neuroanatomical changes such as microcephaly and polymicrogyria [44]. We therefore asked if *grin1^-/-^* fish display gross neuroanatomical changes (**Fig. 1 & Table 3**). We first measured brain morphology at 5 days post-fertilization (dpf) at three different forebrain levels (**Fig. 1A-1F**) (see Materials & Methods) but found no differences (**Figs. 1G-1I**). To determine if morphometric changes might emerge during continued growth, we also measured brain size of live larvae at 6 dpf (**Fig. 1J**) [39] (see Materials & Methods) and brain size relative to body size (**Fig. 1K**), but again found no differences. Finally, we noted qualitatively that the fundamental structure of the *grin1^-/-^* brain, including the overall distribution of cell bodies, formation of subregions, and position of neuropil and fasciculi, was fundamentally unchanged (**Figs. 1A-1F**). Hence, and like murine *Grin1* knockout models [32], the absence of *grin1* in zebrafish does not lead to any notable gross neuroanatomical changes.

**Fig. 1.**
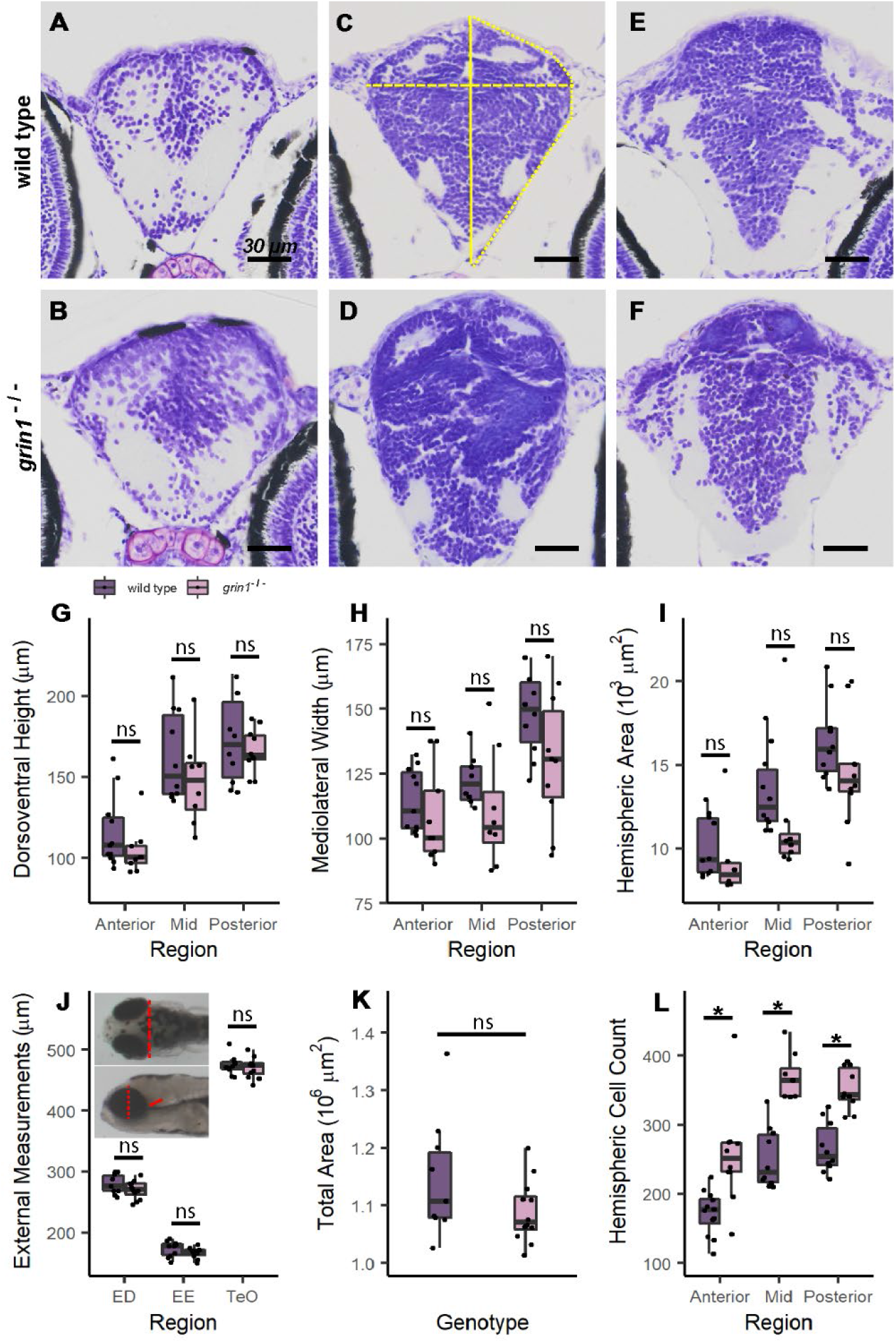
*grin1*^-/-^ fish do not show gross morphological changes in brain structure. **(A**-**F)** Nissl stained 5-micron coronal sections of a 5 dpf wild-type (A, C, & E) and a *grin1*^-/-^ (B, D, & F) fish at the anterior (A & B), mid (C & D) and posterior (E & F) forebrain. Measurement locations indicated in (C): yellow solid line, dorsoventral height measured at midline; dashed line, mediolateral width measured at widest aspect of the brain; and dotted line, hemispheric area. **(G**-**I)** Box and whisker plots indicating median value, interquartile ranges, and maximum and minimum values, for coronal morphometric measurements of dorsoventral height (G), mediolateral width (H) and hemispheric area (I). In this and all subsequent figures, points represent individual values for each fish. **(J & K)** External brain size proxy measurements (J) and total area of fish measured in lateral view (K). Inset to (J): dorsal (*upper*) and lateral (*lower*) views with measurement locations indicated in red dashed line (optic tectum width, TeO), dotted line (eye diameter, ED), and solid line (eye to ear distance, EE). **(l)** Total hemispheric cell counts of 5 dpf fish performed on representative sections at anterior, mid, and posterior levels of the forebrain (A-F) in wild-type and *grin1*^-/-^ fish. Significance (*t-test*) is indicated (*\* p < 0.05; ns*, not significant). For specific sample sizes and statistical values see **Table 3**

**Table 3.** Analysis of gross morphology of wild type and *grin1*^-/-^ fish.

| Metric | Stage | Genotype | n | Mean ± SD<br>($\mu\text{m}$) / ($\mu\text{m}^2$) /<br>(Cell No.) | p value<br>(*t*-test) |
| --- | --- | --- | --- | --- | --- |
| Dorsoventral Height<br>(Anterior) | 5 dpf | wild type | 11 | 120 ± 22 | 0.07 |
| | | *grin1*<sup>-/-</sup> | 9 | 100 ± 15 | |
| Dorsoventral Height<br>(Mid) | “ | wild type | 10 | 160 ± 28 | 0.26 |
| | | *grin1*<sup>-/-</sup> | 8 | 150 ± 27 | |
| Dorsoventral Height<br>(Posterior) | “ | wild type | 10 | 170 ± 28 | 0.48 |
| | | *grin1*<sup>-/-</sup> | 10 | 170 ± 14 | |
| Mediolateral Width<br>(Anterior) | “ | wild type | 11 | 115 ± 12 | 0.41 |
| | | *grin1*<sup>-/-</sup> | 9 | 110 ± 18 | |
| Mediolateral Width<br>(Mid) | “ | wild type | 10 | 120 ± 9.2 | 0.21 |
| | | *grin1*<sup>-/-</sup> | 8 | 110 ± 22 | |
| Mediolateral Width<br>(Posterior) | “ | wild type | 10 | 150 ± 16 | 0.20 |
| | | *grin1*<sup>-/-</sup> | 10 | 130 ± 26 | |
| Hemispheric Area<br>(Anterior) | “ | wild type | 11 | 9800 ± 1900 | 0.26 |
| | | *grin1*<sup>-/-</sup> | 9 | 8700 ± 2300 | |
| Hemispheric Area<br>(Mid) | “ | wild type | 10 | 13400 ± 2400 | 0.27 |
| | | *grin1*<sup>-/-</sup> | 8 | 11600 ± 4000 | |
| Hemispheric Area<br>(Posterior) | “ | wild type | 10 | 16400 ± 2400 | 0.17 |
| | | *grin1*<sup>-/-</sup> | 10 | 14600 ± 3300 | |
| Eye Diameter | 6 dpf | wild type | 10 | 280 ± 16 | 0.16 |
| | | *grin1*<sup>-/-</sup> | 12 | 270 ± 14 | |
| Eye to Ear Distance | “ | wild type | 10 | 170 ± 13 | 0.07 |
| | | *grin1*<sup>-/-</sup> | 12 | 160 ± 11 | |
| Optic Tectum Width | “ | wild type | 10 | 470 ± 16 | 0.55 |
| | | *grin1*<sup>-/-</sup> | 12 | 470 ± 16 | |
| Total Area | “ | wild type | 10 | 1140000 ± 99000 | 0.12 |
| | | *grin1*<sup>-/-</sup> | 12 | 1090000 ± 54000 | |
| Hemispheric Cell<br>Count (Anterior) | 5 dpf | wild type | 12 | 170 ± 32 | 0.006 |
| | | *grin1*<sup>-/-</sup> | 10 | 260 ± 74 | |
| Hemispheric Cell<br>Count (Mid) | “ | wild type | 10 | 250 ± 44 | 9.10 e<sup>-06</sup> |
| | | *grin1*<sup>-/-</sup> | 8 | 370 ± 33 | |
| Hemispheric Cell<br>Count (Posterior) | “ | wild type | 10 | 270 ± 28 | 2.02 e<sup>-05</sup> |
| | | *grin1*<sup>-/-</sup> | 10 | 350 ± 30 | |

Sample size and statistical information for plots in **Fig. 1**.

### *grin1*^-/-^ fish have increased cell densities throughout the forebrain

Despite having comparable gross morphology to wild-type fish, we noted that the brains of *grin1^-/-^* fish were overpopulated with cells (**Fig. 1A-1F**). Initially, to assess this increase, we assayed total hemispheric cell counts at 5 dpf in the anterior, mid, and posterior forebrain, at the same anatomical levels used for morphometric measurements (see Materials & Methods). In each area, we found more cells in the *grin1^-/-^*fish as compared to wild-type fish (**Fig. 1L**).

To begin to define the basis of the increased cell numbers, we carried out a rigorous quantification of cell densities in 9 different regions in the dorsal and ventral telencephalon and diencephalon (**Fig. 2 & Table 4**). By assaying multiple regions, we could assess if there were specific, as opposed to widespread, changes in cell populations. Regions were landmarked anatomically after Mueller & Wullimann (2016) [38], and each was compared at the same representative anatomical level within and between genotypes to ensure equivalent expression boundaries and no double counting of cells (see Materials & Methods).

**Fig. 2.**
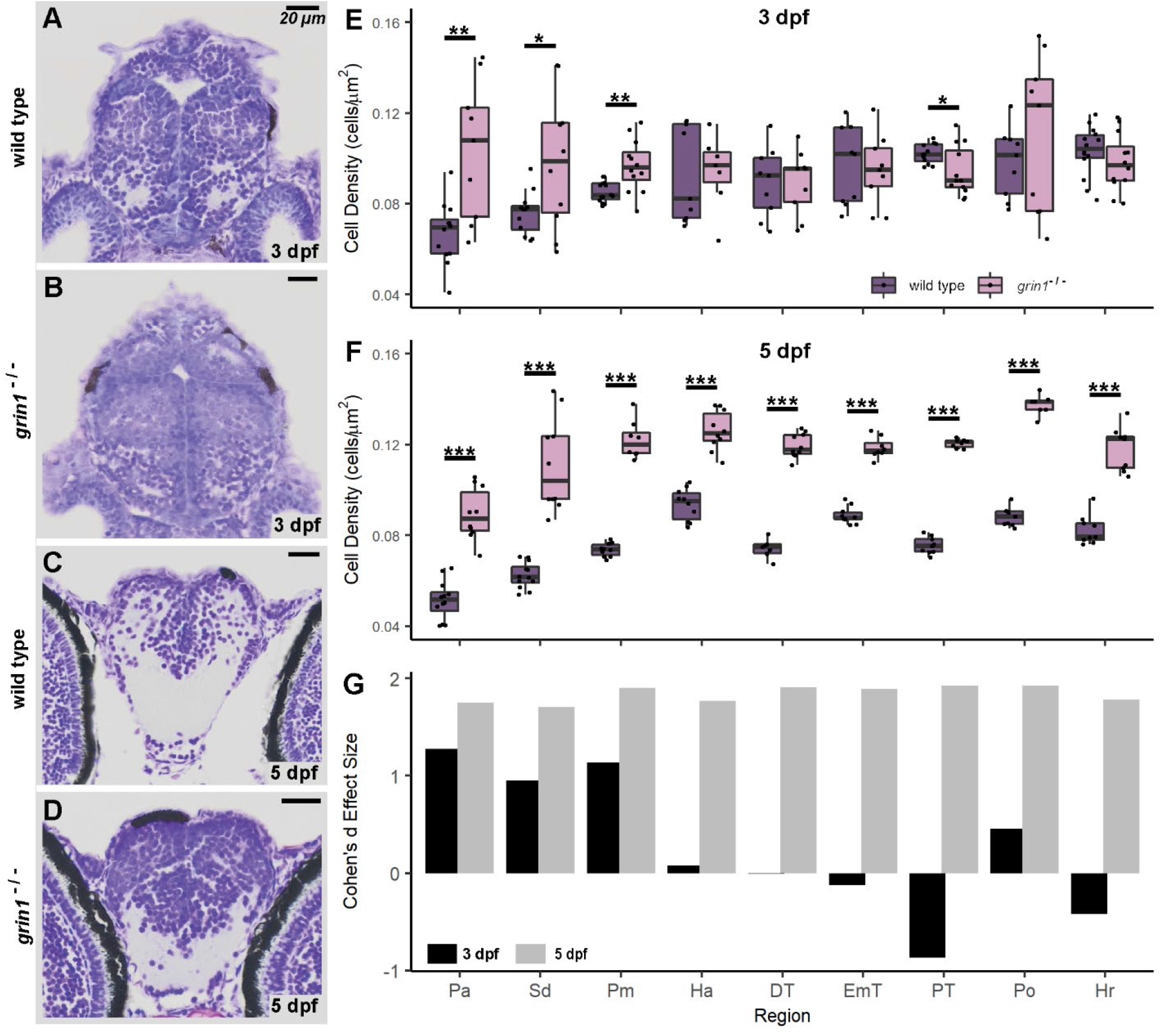
*grin1*^-/-^ fish have increased cell densities throughout the forebrain. **(A**-**D)** Nissl stained 5-micron coronal sections of a 3 dpf wild-type (A) and a *grin1*^-/-^ (B) fish at the precommissural anterior telencephalon, or of a 5 dpf wild-type (C) and a *grin1*^-/-^ (D) fish in the anterior telencephalon at the level of the anterior commissure. (**E & F**) Box and whisker plots of cell density in 9 different anatomical regions in wild-type and *grin1*^-/-^ fish at 3 dpf (E) and 5 dpf (F). **(G)** Data in (E) and (F) expressed as Cohen’s d effect sizes demonstrating magnitude and directionality of changes (positive value indicates greater cell density in *grin1*^-/-^ fish, negative value indicates greater cell density in wild-type fish) in cell density between wild-type and *grin1*^-/-^ fish, at 3 and 5 dpf, in each anatomical region assayed. Significance *(t-test)* is indicated (*\* p < 0.05, **p < 0.01, ***p < 0.001*). For specific sample sizes and statistical values see **Table 4**

**Table 4.**
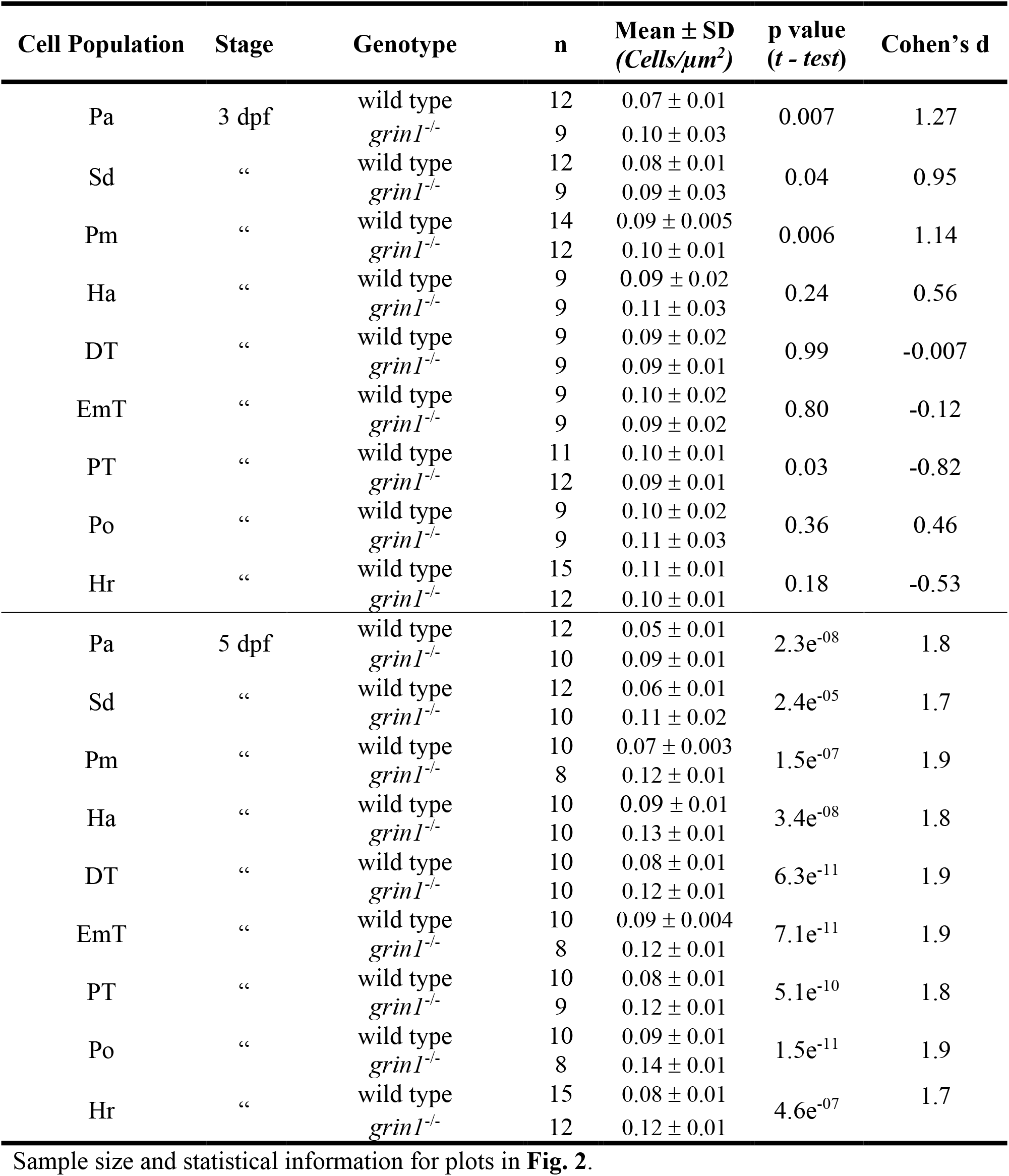
Analysis of forebrain regional cell densities of wild type and *grin1*^-/-^ fish.

| Cell Population | Stage | Genotype | n | Mean ± SD<br>(Cells/$\mu\text{m}^2$) | p value<br>(*t* - test) | Cohen's d |
| --- | --- | --- | --- | --- | --- | --- |
| Pa | 3 dpf | wild type | 12 | 0.07 ± 0.01 | 0.007 | 1.27 |
| | | *grin1*<sup>-/-</sup> | 9 | 0.10 ± 0.03 | | |
| Sd | “ | wild type | 12 | 0.08 ± 0.01 | 0.04 | 0.95 |
| | | *grin1*<sup>-/-</sup> | 9 | 0.09 ± 0.03 | | |
| Pm | “ | wild type | 14 | 0.09 ± 0.005 | 0.006 | 1.14 |
| | | *grin1*<sup>-/-</sup> | 12 | 0.10 ± 0.01 | | |
| Ha | “ | wild type | 9 | 0.09 ± 0.02 | 0.24 | 0.56 |
| | | *grin1*<sup>-/-</sup> | 9 | 0.11 ± 0.03 | | |
| DT | “ | wild type | 9 | 0.09 ± 0.02 | 0.99 | -0.007 |
| | | *grin1*<sup>-/-</sup> | 9 | 0.09 ± 0.01 | | |
| EmT | “ | wild type | 9 | 0.10 ± 0.02 | 0.80 | -0.12 |
| | | *grin1*<sup>-/-</sup> | 9 | 0.09 ± 0.02 | | |
| PT | “ | wild type | 11 | 0.10 ± 0.01 | 0.03 | -0.82 |
| | | *grin1*<sup>-/-</sup> | 12 | 0.09 ± 0.01 | | |
| Po | “ | wild type | 9 | 0.10 ± 0.02 | 0.36 | 0.46 |
| | | *grin1*<sup>-/-</sup> | 9 | 0.11 ± 0.03 | | |
| Hr | “ | wild type | 15 | 0.11 ± 0.01 | 0.18 | -0.53 |
| | | *grin1*<sup>-/-</sup> | 12 | 0.10 ± 0.01 | | |
| Pa | 5 dpf | wild type | 12 | 0.05 ± 0.01 | 2.3e<sup>-08</sup> | 1.8 |
| | | *grin1*<sup>-/-</sup> | 10 | 0.09 ± 0.01 | | |
| Sd | “ | wild type | 12 | 0.06 ± 0.01 | 2.4e<sup>-05</sup> | 1.7 |
| | | *grin1*<sup>-/-</sup> | 10 | 0.11 ± 0.02 | | |
| Pm | “ | wild type | 10 | 0.07 ± 0.003 | 1.5e<sup>-07</sup> | 1.9 |
| | | *grin1*<sup>-/-</sup> | 8 | 0.12 ± 0.01 | | |
| Ha | “ | wild type | 10 | 0.09 ± 0.01 | 3.4e<sup>-08</sup> | 1.8 |
| | | *grin1*<sup>-/-</sup> | 10 | 0.13 ± 0.01 | | |
| DT | “ | wild type | 10 | 0.08 ± 0.01 | 6.3e<sup>-11</sup> | 1.9 |
| | | *grin1*<sup>-/-</sup> | 10 | 0.12 ± 0.01 | | |
| EmT | “ | wild type | 10 | 0.09 ± 0.004 | 7.1e<sup>-11</sup> | 1.9 |
| | | *grin1*<sup>-/-</sup> | 8 | 0.12 ± 0.01 | | |
| PT | “ | wild type | 10 | 0.08 ± 0.01 | 5.1e<sup>-10</sup> | 1.8 |
| | | *grin1*<sup>-/-</sup> | 9 | 0.12 ± 0.01 | | |
| Po | “ | wild type | 10 | 0.09 ± 0.01 | 1.5e<sup>-11</sup> | 1.9 |
| | | *grin1*<sup>-/-</sup> | 8 | 0.14 ± 0.01 | | |
| Hr | “ | wild type | 15 | 0.08 ± 0.01 | 4.6e<sup>-07</sup> | 1.7 |
| | | *grin1*<sup>-/-</sup> | 12 | 0.12 ± 0.01 | | |

Sample size and statistical information for plots in **Fig. 2**.

At the beginning of postembryonic neurogenesis (3 dpf), *grin1^-/-^*fish have greater cell densities in the telencephalon anterior pallium (Pa), dorsal aspect of the subpallium (Sd), and mid pallium (Pm) (**Fig. 2E)**, but in the diencephalon, there was no statistical difference in the habenula (Ha), dorsal thalamus (DT), thalamic eminence (EmT), preoptic area (Po), and rostral hypothalamus (Hr). Interestingly, in one region, the posterior tuberculum (PT), there was a significant decrease in cell density. In contrast, at 5 dpf, which is well into postembryonic neurogenesis, all forebrain populations in the *grin1^-/-^*fish showed significant increases in cell density (**Fig. 2F**).

To compare the magnitude and directionality of the change between genotypes, we calculated Cohen’s d effect size (**Fig. 2G**). At 5 dpf, the effect size was very large (Cohen’s d > 1.2) and comparable across all assayed brain regions, even those that showed no difference (Ha, DT, EmT, Po, Hr) or a reduction (PT) at 3 dpf.

In summary, at the start of postembryonic development, fish lacking NMDA receptors display an excessive number of cells in more rostral forebrain regions with this phenotype extending caudally and becoming widespread during post-embryonic development. This suggests that NMDA receptors are required for proper regulation of cell proliferation.

### NMDA receptors suppress neuron proliferation in the forebrain

The supernumerary cells in the forebrain of *grin1*^-/-^ fish could be neurons or other cell types. To distinguish these alternatives, we assayed expression of Polysialylated-Neural Cell Adhesion Molecule (PSA-NCAM), one of the earliest expressed neural-specific markers [45]. We used PSA-NCAM to capture neuronal identity early in the maturational/differentiation process during this highly proliferative period. We tested expression at 5 dpf, when the cell densities were robustly and uniformly enhanced, and divided our analysis into three gross anatomical regions: precommissural telencephalon, postcommissural telencephalon, and diencephalon (**Fig. 3A-3F**) (See Materials & Methods). Proliferating Cell Nuclear Antigen (PCNA) reactivity was also used to establish an anatomical reference for proliferative zones, at which most cells are PCNA^+^.

**Fig. 3.**
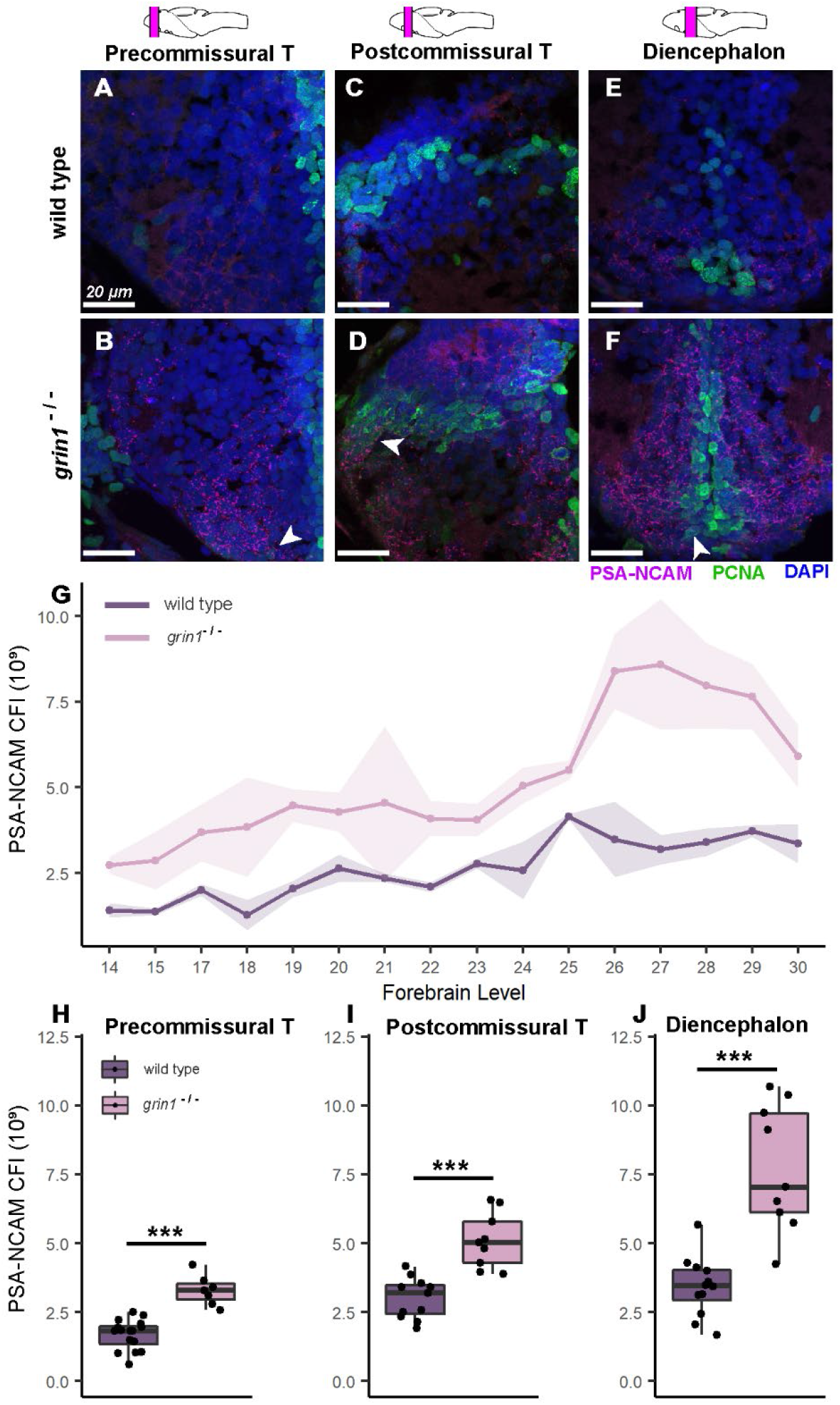
NMDA receptors suppress neuron proliferation in the forebrain. **(A**-**F)** PSA-NCAM (M-Cherry) and PCNA (green) expression with DAPI counterstain in wild-type (A, C, & E) or *grin1*^-/-^ (B, D, & F) fish at 5 dpf either in the precommissural telencephalon (A & B), postcommissural telencephalon (C & D), or diencephalon (E & F). Brain schematics above images indicate the gross anatomical regions sampled. Brain schematics after Mueller & Wullimann (2016). Arrowheads indicate example cells coexpressing PSA-NCAM and PCNA. **(G**) Line graph with points showing the mean PSA-NCAM corrected fluorescence intensity (CFI) at each anatomical level assayed from anterior to posterior (left to right). Ribbon surrounding line graph indicates standard error of the mean. Anatomical levels are numbered according to Mueller & Wullimann (2016) (see Materials & Methods). Wild type n = 16; *grin1*^-/-^ n = 9. *t-test*, *p = 7.3e^-05^*. (*t-test* was performed as a comparison of the aggregate mean CFI across the forebrain for each group.) **(H**-**J)** Box and whisker plots of CFI values within each gross anatomical region. Wild type and *grin1*^-/-^ sample sizes and p value (*t-test*): Precommissural telencephalon (H), 16, 7; *p = 1.7e-^06^*. Postcommissural telencephalon (I), 11, 9; *p = 4.4e^-05^*. Diencephalon (J), 12, 9; *p = 0.0003.* Significance (*t-test*) is indicated (*\*\*\*p < 0.001*)

To obtain an overview of PSA-NCAM expression, we performed corrected fluorescence intensity analysis (CFI) on each atlas level (See Materials & Methods). First, we determined average expression for each genotype at multiple anatomical levels across the forebrain (**Fig. 3G**) and found elevated expression of PSA-NCAM in *grin1^-/-^* fish at every level. We then assessed differential expression in each gross anatomical region of the forebrain by averaging the CFI for all atlas levels within each region for each individual specimen. In this way, we obtained an average PSA-NCAM expression in each of three forebrain regions for each fish. We found that in every gross anatomical region, *grin1^-/-^*fish have significantly more PSA-NCAM expression than their wild-type counterparts (**Fig. 3H, 3I, & 3J**). These data indicate that the increased cell densities observed in *grin1^-/-^* fish reflect an increase in the number of forebrain neurons.

### Supernumerary neurons in the forebrain of *grin1*^-/-^ fish do not result from a failure of programmed cell death

Programmed cell death is a natural part of neurodevelopment [46]. The supernumerary immature neuron population observed in *grin1^-/-^* fish could therefore result from failure of programmed cell death. To test this possibility, we assayed activated caspase-3, a death protease characteristic of apoptosis [47], on wholemount zebrafish larvae at 3 dpf and found no difference in activated caspase-3 expression between wild-type and *grin1^-/-^* fish (**Fig. S1**). This result suggests that the supernumerary neurons in the mutant brains do not arise from decreased cell death but rather increased neurogenesis.

### Radial glia are not increased in the forebrain of *grin1^-/-^* fish

The terms used in neurogenesis are variable between experimental models and scientific specialties. For clarity, we use the term neural stem cell (NSC) to specifically refer to RGCs. We define the term neuroblast as the neural-fated progeny of a NSC that is not yet fully differentiated and an NPC as a neuroblast that is immature and actively mitotic. In mammals, NPCs encompass transit amplifying cells (tAC), and their subtype, intermediate progenitor cells [48]. The term intermediate progenitor cell is not used in zebrafish for several reasons including structural differences in the germinative zones. Hence, we use tACs to define all NPCs in zebrafish. Zebrafish also have NSC populations composed of neuroepithelial cells that persist beyond early neurodevelopment [49]. These populations are limited in the forebrain and primarily supply neurons to regions that were not assayed, such as the olfactory bulb, and thus were excluded from our study.

The PSA-NCAM results (**Fig. 3**) indicate a preponderance of immature neurons. These excessive neurons could arise directly from increases in the population and/or proliferation of NSCs, or indirectly from excessive proliferation of tACs. Glial Fibrillary Acidic Protein (GFAP) is robustly expressed in RGCs, the principal NSCs of the forebrain [50,51]. Hence, to determine if increased NSC populations were responsible for the supernumerary neurons seen globally at 5 dpf, we assayed GFAP expression at 3 dpf (**Fig. 4**). GFAP is primarily expressed in the processes of RGCs, therefore, to capture the full extent of expression, we performed CFI analysis (see Materials & Methods). In the three gross anatomical regions assayed – the precommissural telencephalon (**Fig. 4A & 4B**), postcommissural telencephalon (**Fig. 4C & 4D**), and diencephalon (**Fig. 4E & 4F**) – we found no difference in GFAP expression between wild-type and *grin1^-/-^*fish (**Fig. 4G-4I**). Thus, in the forebrain regions analyzed, the absence of NMDA receptors does not lead to enhanced numbers of RGCs.

**Fig 4.**
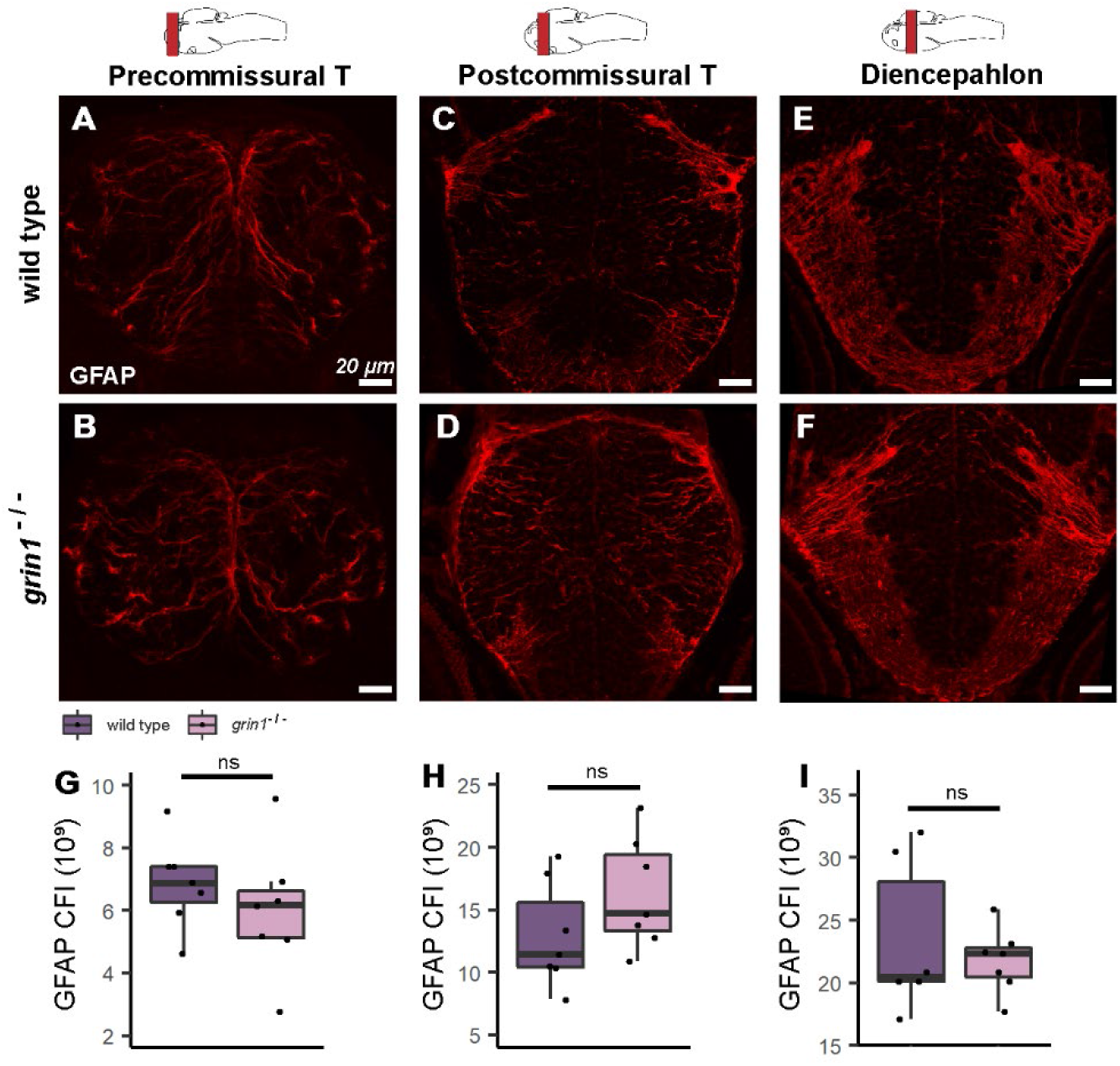
Radial glia are not increased in the forebrain of *grin1^-/-^* fish. **(A**-**F)** GFAP expression (index of RGCs) of 3 dpf wild-type (A, C, & E) and *grin1*^-/-^ (B, D, & F) fish at three gross anatomical levels of the forebrain (see Materials & Methods). Brain schematics above images indicate the gross anatomical regions sampled. Brain schematics after Mueller & Wullimann (2016). **(G**-**I)** Box and whisker plots of CFI analysis. Wild type and *grin1*^-/-^ sample sizes and p value (*t-test*): Precommissural telencephalon (G), 7, 7; *p = 0.38*. Postcommissural telencephalon (H), 7, 7; *p = 0.17*. Diencephalon (I), 6, 7; *p = 0.53*. Significance (*t-test*) is indicated (*ns*, not significant)

### NMDA receptors suppress proliferation in neuroblasts (tAC)

The absence of NMDA receptors leads to an enhanced immature neuron population (**Fig. 3**). The source of these extra neurons remains unknown but does not reflect enhanced NSC populations (**Fig. 4**). Nevertheless, the balance between quiescence and proliferation in RGCs could be shifted toward proliferation thus expanding the neuron population. Alternatively, or in addition, neuroblasts could be the source of the observed dysregulation. Neuroblasts can mature into post-mitotic neurons promptly after birth or act as tACs by progressing through one or more rounds of mitosis before maturing; thus, a change to this amplifying phase could cause or contribute to supernumerary neurons. To distinguish these alternatives, we assayed PCNA, which detects both proliferating RGCs and tACs, and took advantage of the different anatomical location and morphology of RGCs and tAC to distinguish these populations (**Fig. 5**) [52].

**Fig. 5.**
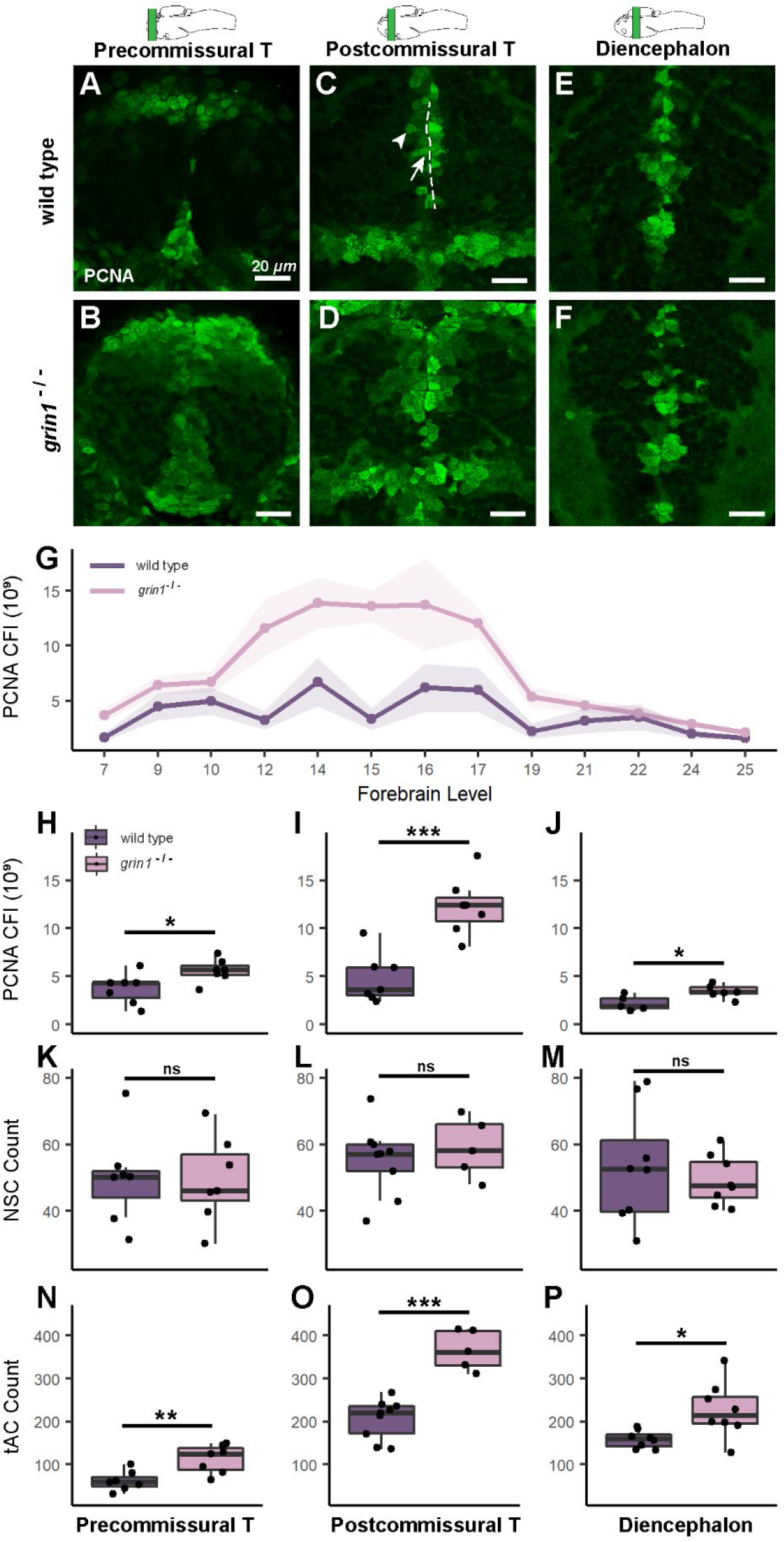
NMDA receptors suppress proliferation in neuroblasts (transit amplifying cells) **(A**-**F)** PCNA expression in wild-type (A, C, & E) and *grin1*^-/-^ (B, D, & F) fish at 3 dpf in the precommissural telencephalon (A & B), postcommissural telencephalon (C & D) and diencephalon (E & F). PCNA is enriched in abventicular and cell rich regions of the parenchyma in the mutant fish. In (C), sample of PVZ (dotted line), example RGC at PVZ (arrow), example abventricular tAC (arrowhead). Brain schematics above images indicate the gross anatomical regions sampled. Brain schematics after Mueller & Wullimann (2016). **(G**) Line graph with points showing the mean corrected fluorescence intensity (CFI) for each genotype, at each anatomical level assayed from anterior to posterior (left to right). Ribbon surrounding line graph indicates standard error of the mean. Anatomical levels are numbered according to Mueller & Wullimann (2016) (see Materials & Methods). Wild type n = 7; *grin1*^-/-^ n = 7. *t-test*, *p = 0.0009*. **(H**-**P)** Box and whisker plots of different analysis of three anatomical levels: precommissural telencephalon, postcommissural telencephalon, and diencephalon. Wild type and *grin1*^-/-^ sample sizes and p value (*t-test*): (**H**-**J)** PCNA CFI values: precommissural telencephalon (H) 7, 7, *p = 0.03*; postcommissural telencephalon (I) 7, 7, *p = 0.0003*; diencephalon (J) 5, 7, *p = 0.01*. **(K**-**M)** PCNA^+^ NSC counts: precommissural telencephalon (K) 7, 7, *p = 0.95*; postcommissural telencephalon (L) 9, 5, *p = 0.54*; diencephalon (M) 8, 8, *p = 0.54*. **(N**-**P**) PCNA^+^ tACs cell counts: precommissural telencephalon (N) 7, 7, *p = 0.005*; postcommissural telencephalon (O) 9, 5, *p = 5.1e^-05^*; diencephalon (P) 8, 8, *p = 0.02*. Significance (*t-test*) is indicated (*\* p < 0.05, **p < 0.01, ***p < 0.001; ns, not significant*)

We assayed PCNA expression in three gross anatomical regions (**Fig. 5A-5F**) (see Materials & Methods). PCNA expression in *grin1^-/-^* fish was enhanced across the forebrain (**Fig. 5G**) and within each specific gross brain region (**Fig. 5H-5J**) with the greatest difference between genotypes in the postcommissural telencephalon (**Fig. 5I)**.

CFI provides regional, not cellular resolution. To address whether this enhanced PCNA activity arose from RGCs or tACs, we quantified proliferating RGC and tAC populations (**Fig. 5K-5P**). NSCs and tACs were identified by location (PVZ, apical location for NSCs and abventricular, non-apical for tACs) (**Fig. 5C**), somal shape (ovoid for RGCs and spherical for tACs) [53,52], and GFAP reactivity, which was used as a distinguishing factor for RGCs in conjunction with location and morphology to differentiate these two NSPC populations despite the mild persistence of GFAP expression in newborn neuroblasts (**Fig. S2**) [54]. In all forebrain regions, we found no difference in the number of PCNA^+^ RGCs (**Fig. 5K-5M**) but found significantly more PCNA^+^ tACs, with the largest difference in the postcommissural telencephalon (**Fig. 5N-5P**).

Together, these data indicate that without NMDA receptors, the RGC proliferation is unchanged, but that there is an increase in the number of proliferating tACs. Furthermore, since neither the RGC population (**Fig. 4**) nor their proliferation (**Fig. 5K-5M**) is increasing, the RGCs are presumably not producing more tACs than in wild type. This suggests that in the *grin1*^-/-^ fish, the tACs are, themselves, the source of their own enriched population. Enhanced populations of tACs, combined with their abnormally prolonged mitotic competency, leads to escalation of supernumerary neurons during postembryonic development. Consistent with this idea, we observed a subset of PSA-NCAM^+^ cells that were also PCNA^+^ (**Fig. 3B, 3D, & 3F, arrowheads**), indicating that in the absence of NMDA receptors the tACs are still mitotic after the onset of neuronal marker expression.

### Transit amplifying cells express GluN1

Our findings suggest that NMDA receptors are required for limiting tAC proliferation. A key question is whether these cells normally express NMDA receptors. If so, this would suggest that the activation of NMDA receptors suppresses mitosis in tACs and caps neurogenic activity, thus explaining the regulatory loss and supernumerary neurons in *grin1*^-/-^ fish. To address this question, we performed IHC in wild-type fish at 3 dpf using antibodies against GluN1 and PCNA to test whether tACs, which express PCNA, coexpress GluN1 (**Fig. 6**). In these fish, GluN1 was expressed robustly on cell membranes and in the neuropil (**Fig. 6A & 6B**), consistent with the expected expression on the plasma membrane and in the dendrites. Further, PCNA^+^ cells in abventricular and cell-rich locations (**Fig. 6C & 6D**) also coexpress GluN1 on their surrounding plasma membrane (**Fig. 6A-6F, arrows**). Thus, proliferating tACs express NMDA receptors on the plasma membrane.

**Fig. 6.**
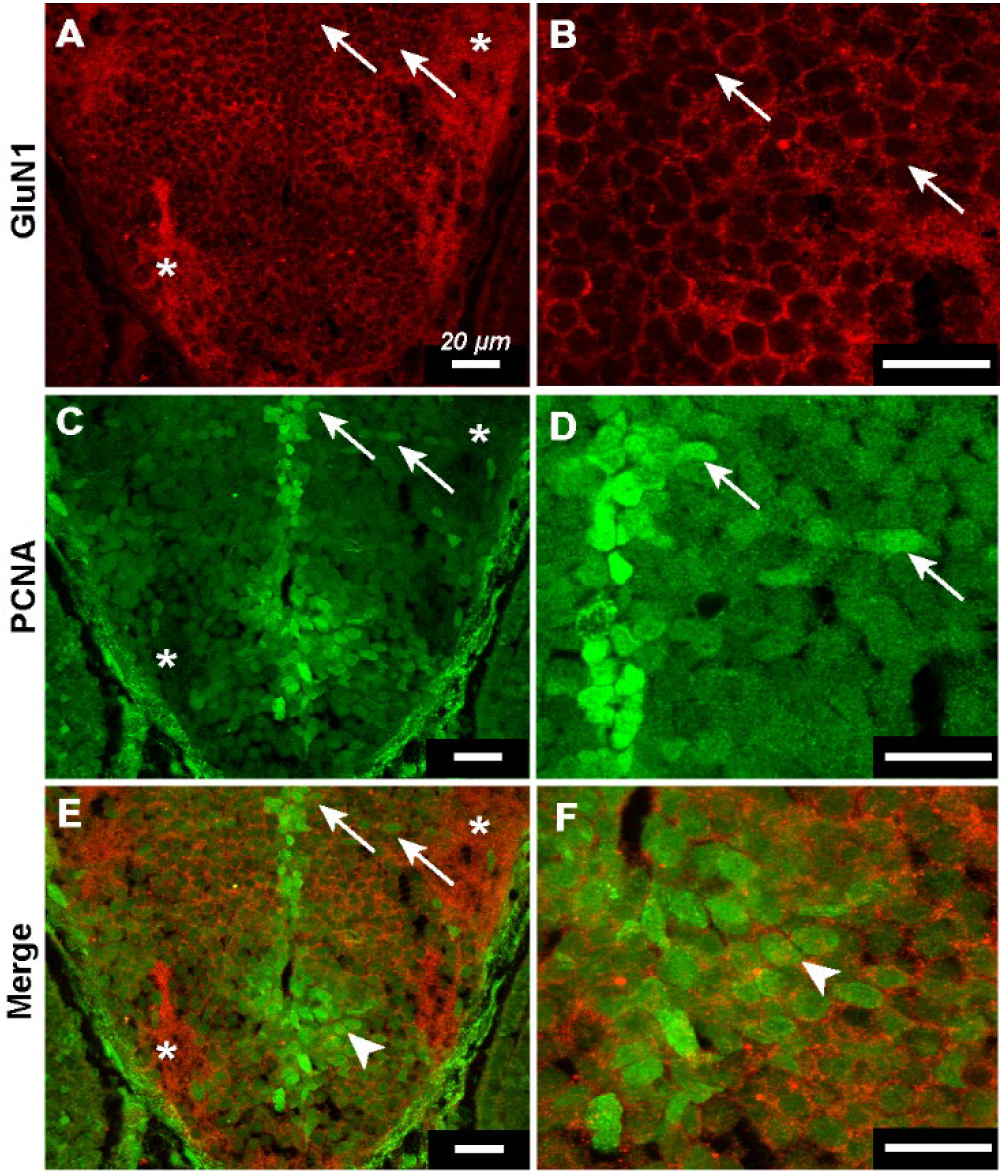
Transit amplifying cells express GluN1. **(A**-**F)** Immunohistochemistry against GluN1 and PCNA in 3 dpf wild-type fish. Panels on the right are magnified versions of those on the left. Asterisks indicate example regions of neuropil in (A, C, & E). **(A & B)** GluN1 channel showing robust plasmelemmal and neuropilar dendritic expression of GluN1. Arrows indicate example cells in abventicular and cell-rich parenchymal locations (A) and illustrate plasmelemmal GluN1 expression (B). **(C & D)** PCNA channel showing ventricular PCNA^+^ NSCs and abventricular and cell-rich parenchymal PCNA^+^ tACs cells. Arrows indicate the same cells as in (A & B) and illustrate that the same cells express both PCNA and GluN1. **(E & F)** Merged GluN1 and PCNA channels. Arrows indicate same cells as in (A, B, C, & D) arrow heads indicate example PCNA^+^ tAC with GluN1 puncta.

### KCC2 and PCNA expression are inversely correlated in the forebrain of wild-type fish

The absence of NMDA receptors results in an excessive number of proliferating tACs (**Fig. 5**) that inappropriately amplify the neuronal pool (**Fig. 3**). Why there is an excessive number is unknown but does not appear to be dependent on the source of tAC as we observed no change in RGC populations (**Fig. 4**) or mitotic activity (**Fig. 5**). An alternative explanation is that in the absence of NMDA receptor-mediated signaling, neuroblasts acting as tACs no longer mature and differentiate into postmitotic neurons in an appropriate, timely fashion. To initially address this question, we studied the expression of the chloride transporter, KCC2, which is one of the earliest markers of neuronal maturation [22]. Still, little work has been done on KCC2 in zebrafish; therefore, we wanted to validate its expression in wild-type fish (**Fig. 7**). Additionally, though KCC2 upregulation is associated with GABA’s inhibitory actions on mature neurons, the excitatory effect of GABA during development has been linked to both promotion and suppression of mitosis depending on the progenitor cell type [55–57]. Thus, we sought to validate that KCC2 upregulation was correlated with cessation of mitotic activity in the maturing neuroblast.

**Fig. 7.**
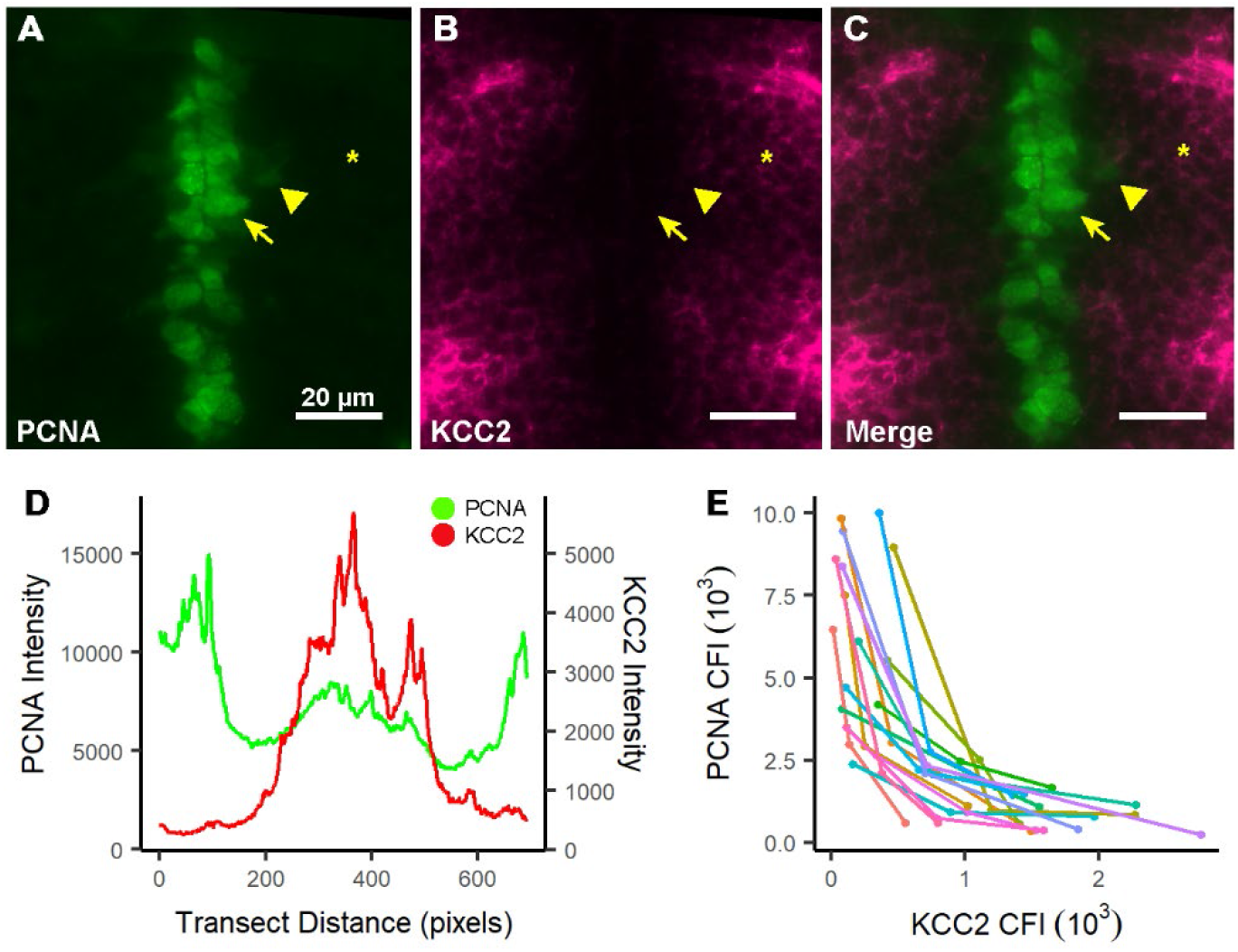
KCC2 and PCNA expression are inversely correlated in the forebrain of wild-type fish. **(A**-**C)** A representative section from the diencephalon of 3 dpf fish showing PCNA (A) and KCC2 (B) expression, and merged image (C). Symbols indicate the same cells in each image: arrows, PCNA^+^ cell with high expression near the PVZ; arrowhead, a PCNA^+^ cell with moderate to low expression at an abventricular position; and asterisk, a PCNA^-^ cell in a perineuropilar position. These cells have no, moderate, and high KCC2 expression, respectively. **(D)** Line graph of corrected line fluorescence (CLF) for PCNA (green) and KCC2 (red) from a representative transect in the telencephalon, showing the inverse expression relationship between PCNA and KCC2. The transect was drawn from the midline PVZ to the tela choroidea at the outermost aspect of the brain. **(E**) Scatter plot showing the inverse relationship between PCNA and KCC2 expression in individual wild-type cells. Each set of 3 color coded dots with their connecting line represents a triplet of cells from one specimen. For each sampling, cell 1 is within the abventricular zone with high PCNA expression (A, B, & C, arrow), cell 2 is within the cell-rich parasagittal zone with moderate PCNA expression (A, B, & C, arrowhead), and cell 3 is in the peri-neuropilar zone with low PCNA expression (A, B, & C, asterisk). Scatter indicates the PCNA and KCC2 CFI value within each individual cell represented as PCNA vs. KCC2. n = 16.

Initially, we analyzed the line fluorescence for KCC2 and PCNA along transects through the dorsal telencephalon of 3 dpf wild-type fish (**Fig. 7A-7D**) (see Materials & Methods) and found an inverse relationship between KCC2 and PCNA that exists across the entire hemisphere in the telencephalon (**Fig. 7D**). Next, we sampled sets of three individual cells in the diencephalon of wild type fish, at increasing migratory distance from the proliferative zone (abventricular, parasagittal, and perineuropilar) (see Materials & Methods). We analyzed the relative coexpression of KCC2 and PCNA of each cell using CFI analysis (**Fig. 7A, 7B, 7C, & 7E**) and found an inverse relationship within each triplet of cells (**Fig. 7E**). This is represented in the traces of cell triplets, 1 from each region of migration from the PVZ, showing that along the migratory path cells with a higher PCNA expression have lower KCC2 expression and vice versa (**Fig. 7A, 7B, 7C, & 7E**).

In summary, these data indicate that KCC2 upregulation is correlated with the maturational cessation of mitosis in tACs and can be used as an index of cell maturation.

### Loss of NMDA receptor function results in reduced KCC2 expression

To determine the maturation status of neuroblasts, we analyzed the KCC2 expression in 3 dpf *grin1^-/-^*fish as compared to wild-type fish (**Fig. 8**). Because neuroblasts begin the maturational process as they migrate [58–60], we performed a line fluorescence analysis in the diencephalon, where the orientation of RGC fibers, and thus the path of migrating neurons, can be reliably interpreted. We analyzed a transect through the dorsal thalamus and one through the posterior tuberculum (**Fig. 8A & 8B**) (see Materials & Methods). We found that *grin1^-/-^* fish have lower maximum average expression of line fluorescence and lower intermittent peaks than wild type fish in both dorsal (**Fig. 8C**) and ventral (**Fig. 8D**) transects.

**Fig. 8.**
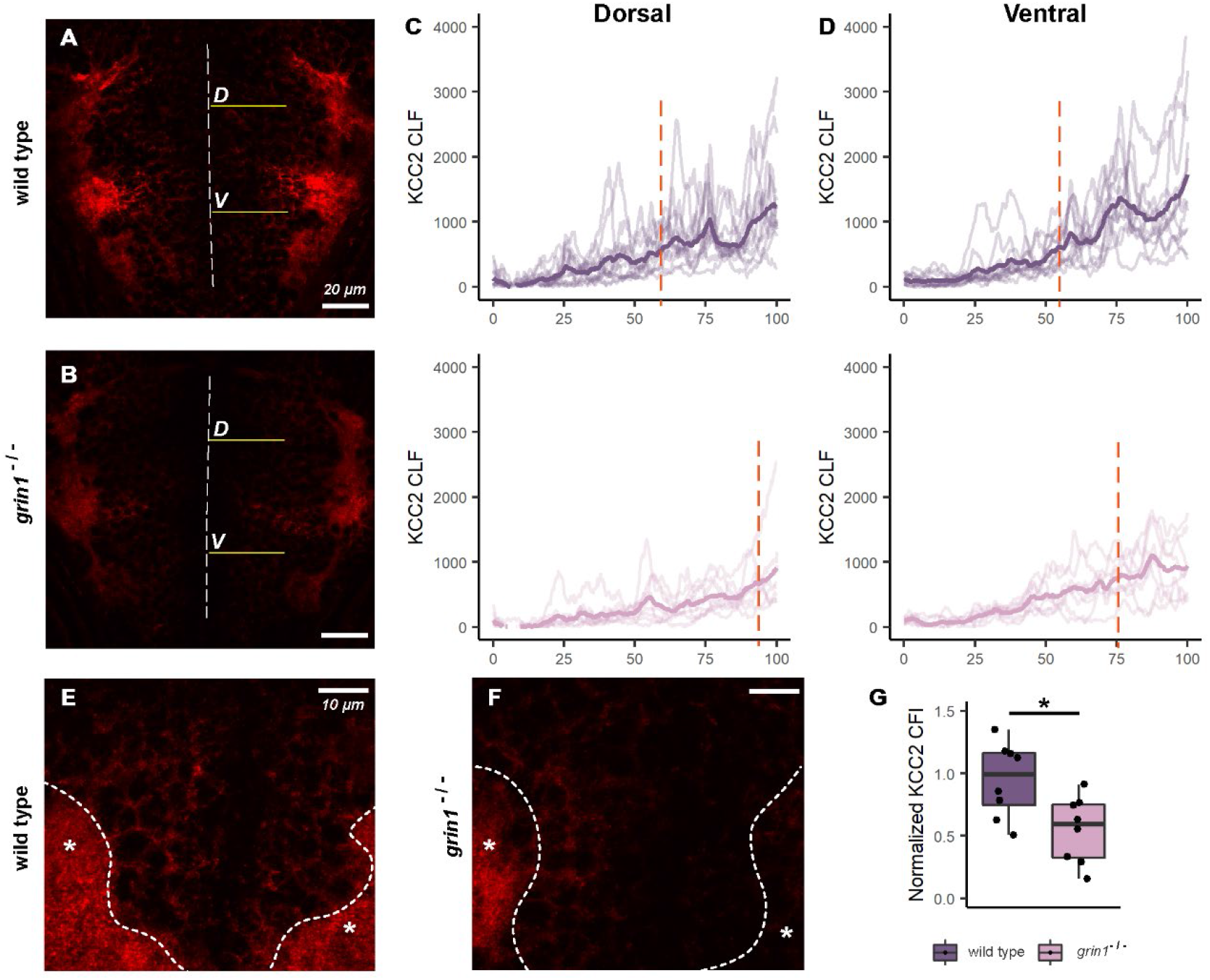
Loss of NMDA receptor function results in reduced KCC2 expression. **(A & B)** Representative images of the diencephalon of a wild-type (A) and *grin1*^-/-^ (B) fish. Dashed line is the midline proliferative zone, whereas the solid lines represent the transects drawn for corrected line fluorescence intensity analysis (CLFI). The dorsal transect (labeled D) passes through the dorsal thalamus, while the ventral transect (labeled V) passes through the posterior tuberculum. **(C** & **D)**. Line graphs showing CLFI for dorsal (C) and ventral (D) transects in wild-type (top) and *grin1*^-/-^ (bottom) fish. Low opacity lines represent individual traces. Bold lines represent the averages of all traces binned in increments of 0.5% along the transect length. Peaks represent points at which the transect crosses the plasma membrane. The dotted orange line represents the distance along the transect at which the average intensity reaches half the wild-type average maximum value. **(E & F)** Representative images of the cell-rich region of the anterior telencephalon in wild-type (E) and *grin1*^-/-^ (F). Asterisks indicate regions of the anterior commissure in each image and the area above the dashed line indicates the cell-rich region surrounding the anterior commissure. **(G)** Box and whisker plot of normalized CFI for the cell rich region surrounding the anterior commissure. Wild type n = 8, *grin1*^-/-^ n = 8, *t-test p = 0.01*. Significance (*t-test*) is indicated (*\*p < 0.05*)

Neuroblast maturation, as evidenced by KCC2 expression, is a continuum. Maximal levels of KCC2 expression are not required for a neuron to begin to mature [61]. Thus, we assessed the distance from the proliferative zone at which we first see 50% of the maximum average wild-type expression of KCC2 as a measure of initiation of the maturational process. The wild type dorsal transect average reaches half maximum at approximately 60% of the transect length, whereas in *grin1*^-/-^ fish, the transect average reaches half maximum at approximately 90% of the transect length (**Fig. 8C**). Similarly in the ventral transect, the wild-type average reaches half maximum at approximately 58% of the transect distance, whereas *grin1*^-/-^ does not reach half maximum until more than 75% of the transect distance (**Fig. 8D**).

To assess total cell expression of KCC2, we performed CFI analysis on the cell-rich compartment at the level of the anterior commissure (**Fig. 8E-8G**). KCC2 is also robustly expressed on dendrites. Therefore, we used the level of the anterior commissure to exclude the dendritic compartment from this analysis to avoid potential confounds of NMDA receptor effects on arborization [13,62] (see Materials & Methods). Furthermore, as the amount of KCC2 expression must be considered relative to the total number of cells, we normalized the KCC2 CFI against the amount of DAPI^+^ cells. We find that relative to wild type, *grin1^-/-^* fish have significantly less KCC2 expression in the cell rich portion of the forebrain (**Fig. 8G**). Notably, though excluded from this analysis, reduced expression of KCC2 in the dendritic compartment is visually apparent in *grin1*^-/-^ fish as well as in the cell-rich region (**Fig. 8F**).

In summary, these data, taken together, demonstrate a lower overall expression of KCC2 paired with a delay in expression as measured by distance from the proliferative zone of the neuroblast. Hence, without NMDA receptors, there is a delay in the maturation of tACs.

### NMDA receptor ionotropic function is required for suppression of neurogenesis

NMDA receptors act via multiple signaling mechanisms including Ca^2+^ influx and non-ionotropic activity [12,63] that could be responsible for the *grin1*^-/-^ phenotype. To distinguish between these alternatives, we used the non-competitive inhibitor MK-801, which blocks the ionotropic signaling of NMDA receptors, including Ca^2+^ influx, but not non-ionotropic signaling [64,63]. Beginning at 24 hours post fertilization, which is the earliest observed expression of NMDA receptors in zebrafish [65,34], wild-type fish were treated daily with 40 µM MK-801. As observed in *grin1*^-/-^ fish (**Fig. 3**), MK-801 in wild-type fish causes increased cell densities in most forebrain regions (Pa, Sd, Pm, DT, PT, and Po) by 5 dpf (**Fig. 9 & Table 5**). Recapitulation of the *grin1^-/-^* phenotype was not observed in every region (EmT and Hr) which may reflect limitations of pharmacological blockade. Nevertheless, our treatment of wild-type fish largely reproduced the *grin1*^-/-^ phenotype. Thus, ionotropic signaling through NMDA receptors is necessary for the maturation and termination of proliferation in tACs.

**Fig. 9:**
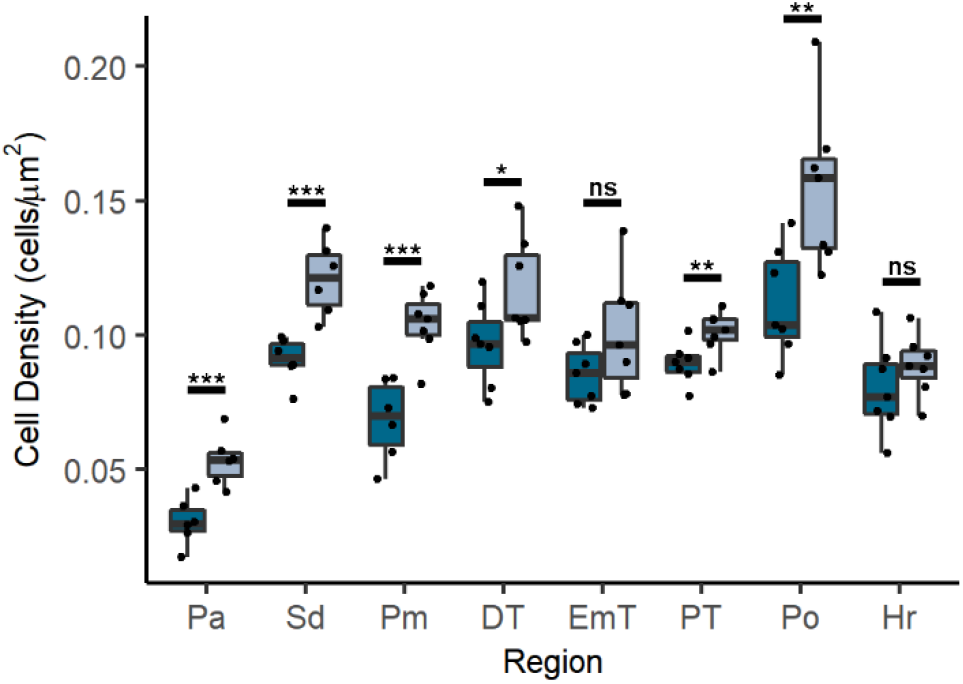
NMDA receptor ionotropic function is required suppression of neurogenesis. Box and whisker plot of cell density in 8 anatomical regions in control and MK-801 treated fish at 5 dpf. Significance *(t-test)* is indicated (*\* p < 0.05, **p < 0.01, ***p < 0.001*; *ns*, not significant). For specific sample sizes and statistical values see **Table 5**

**Table 5.**
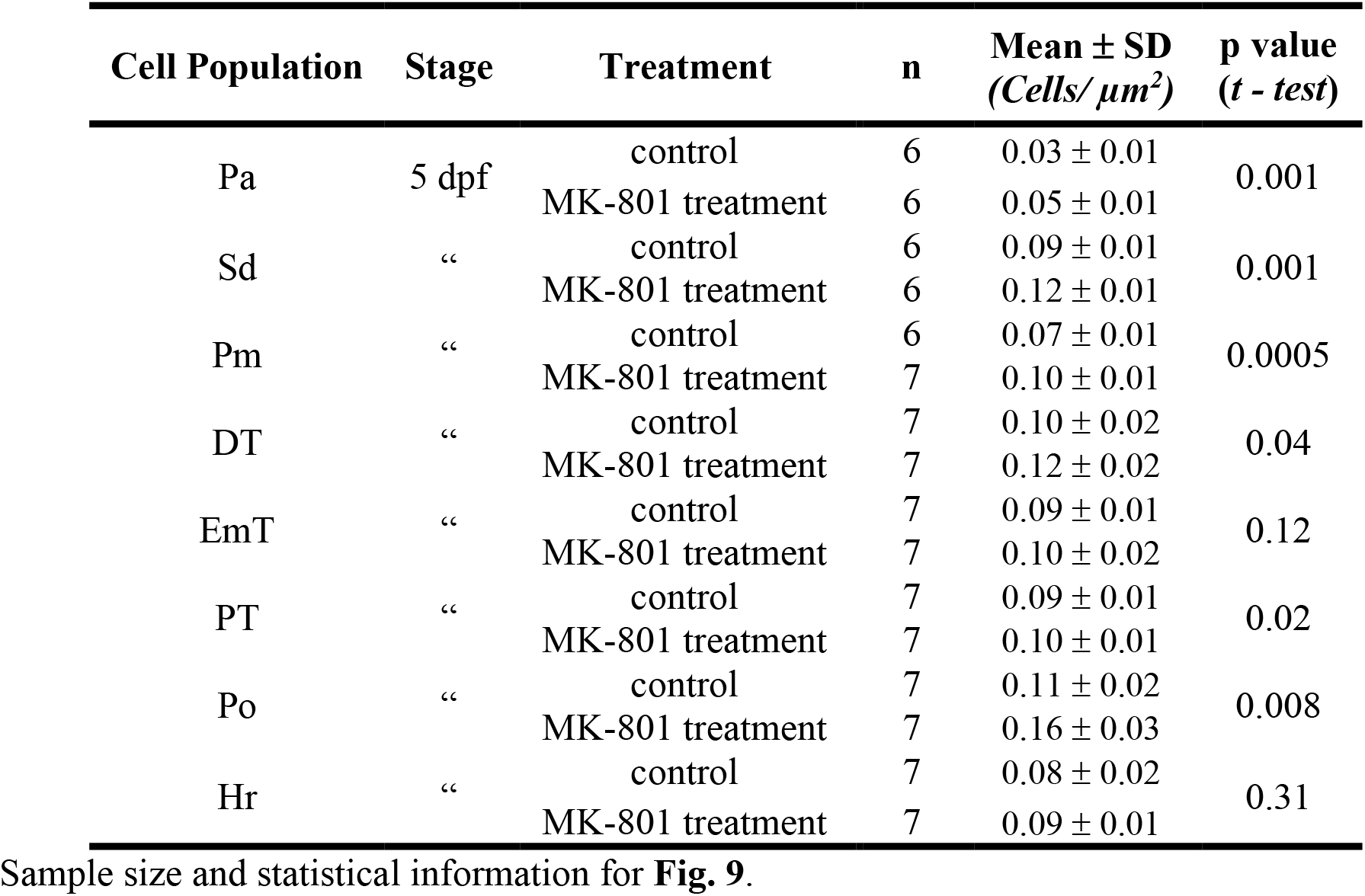
Statistical data for regional cell densities in control and MK-801 treatment fish.

| **Cell Population** | **Stage** | **Treatment** | **n** | **Mean ± SD**<br>**(Cells/$\mu\text{m}^2$)** | **p value**<br>**(t - test)** |
| --- | --- | --- | --- | --- | --- |
| Pa | 5 dpf | control | 6 | $0.03 \pm 0.01$ | 0.001 |
| | | MK-801 treatment | 6 | $0.05 \pm 0.01$ | |
| Sd | “ | control | 6 | $0.09 \pm 0.01$ | 0.001 |
| | | MK-801 treatment | 6 | $0.12 \pm 0.01$ | |
| Pm | “ | control | 6 | $0.07 \pm 0.01$ | 0.0005 |
| | | MK-801 treatment | 7 | $0.10 \pm 0.01$ | |
| DT | “ | control | 7 | $0.10 \pm 0.02$ | 0.04 |
| | | MK-801 treatment | 7 | $0.12 \pm 0.02$ | |
| EmT | “ | control | 7 | $0.09 \pm 0.01$ | 0.12 |
| | | MK-801 treatment | 7 | $0.10 \pm 0.02$ | |
| PT | “ | control | 7 | $0.09 \pm 0.01$ | 0.02 |
| | | MK-801 treatment | 7 | $0.10 \pm 0.01$ | |
| Po | “ | control | 7 | $0.11 \pm 0.02$ | 0.008 |
| | | MK-801 treatment | 7 | $0.16 \pm 0.03$ | |
| Hr | “ | control | 7 | $0.08 \pm 0.02$ | 0.31 |
| | | MK-801 treatment | 7 | $0.09 \pm 0.01$ | |

Sample size and statistical information for **Fig. 9**.

## DISCUSSION

NMDA receptors are expressed in NSPCs [19,20] and are associated with neurodevelopmental disorders, but their contribution to neurogenesis is not fully understood. Here, we demonstrate that NMDA receptors are required for the suppression of indirect neurogenesis. We find that without NMDA receptor-mediated signaling, supernumerary neurons develop in the forebrain (**Fig. 2 & 3**). This effect begins during embryonic development in the anterior forebrain, producing mild and regionally constrained phenotypes by 3 dpf (**Fig. 2E**), but escalates during postembryonic neurogenesis to produce very large effect sizes (Cohen’s d > 1.2) in all forebrain regions assayed by 5 dpf (**Fig. 2F & G**). This increased neuronal density does not arise directly from the NSC pool (**Figs. 4 & 5K-M**) but rather indirectly, from neuroblasts that are mitotically active (tACs) (**Fig. 5N-P**). We demonstrate that with loss of NMDA receptor function, neuroblasts acting as tACs do not mature into post mitotic neurons in a timely manner as demonstrated by reduced KCC2 expression (**Fig. 8**). This prolongs their mitotic stage and inappropriately amplifies the neuronal pool.

In zebrafish, all mitotically-active neuroblasts, or NPCs, are referred to as transit amplifying cells (tAC). In mammals, an important subset of NPCs of the cortex are referred to as intermediate progenitor cells (iPCs). Despite the lack of this terminology in zebrafish, some of the cells we term tACs are akin to iPCs as they express Tbr2 [66] and are generated in the anatomical regions, such as the dorsal pallium, that rely on these progenitors for generation of cortical projection neurons [67]. Nevertheless, we discovered supernumerary neurons in all regions of the forebrain, even those homologous to regions that do not generate neurons from cells strictly described as iPCs. Thus, NMDA receptors are required for suppression of neurogenesis in all subtypes of amplifying, neuronally restricted cells.

NMDA receptors are well-known to mediate synaptic excitatory neurotransmission. In NSPCs, there are no synapses so any signaling via NMDA receptors must occur via non-synaptic mechanisms (paracrine or autocrine) [68]. Non-synaptic neurotransmitter signaling sources have been attributed to NSPCs [69], Cajal-Retzius cells [70], cerebral spinal fluid [71], as well as endothelial cells of blood vessels [72]. Non-synaptic signaling can be either ionotropic or non-ionotropic. Nevertheless, the excessive proliferation of neuroblasts persists in wild-type fish treated with MK-801, an NMDA receptor channel blocker (**Fig. 9**). The maturation of tACs is therefore dependent on ionotropic signaling, most likely Ca^2+^ influx.

### The spatiotemporal trajectory of developmental neurogenesis

Although we focused on the postembryonic period, we observed supernumerary neurons present in the anterior forebrain at 3 dpf (**Fig. 2E**). As 3 dpf marks the beginning of postembryonic development in the zebrafish, this is a process that likely began during embryonic development and then progressed in magnitude and extent during the postembryonic period. Nevertheless, in the more posterior areas of the forebrain, the diencephalic populations, we saw different degrees of neurogenesis suppression at 3 dpf, indicated by a negative Cohen’s d effect size in most regions (**Fig. 2G**). Hence, at early stages in different subsets of NSPCs, NMDA receptors may have different roles.

During early neurodevelopment in the zebrafish, formation of the neural tube, regional subdivisions, and ensuing rounds of neurogenesis generally progress along the anterior to posterior axis [73]. This spatiotemporal character of neurogenesis means that different stages of brain architecture are unfolding in different regions at different times and, thus, will result in different findings in different regions at any temporal window. Our PCNA analysis of proliferation at 3 dpf (**Fig. 5**) may reflect these temporal differences: the more anterior precommissural telencephalon shows a mild increase, the postcommissural telencephalon the greatest increase, and the diencephalon again a weak increase. Paired with the cell density data at 3 dpf (**Fig. 2E**), one interpretation of these data is that at 3 dpf we observed a dysregulatory process that has already partially unfolded in the anterior portion of the forebrain, is actively underway in the mid forebrain, and has not yet begun in the posterior forebrain. Similarly, the PSA-NCAM analysis of immature neurons at 5 dpf (**Fig. 3**) reveals the greatest increase in the diencephalon (**Fig. 3G**). That, paired with the cell density phenotype, which progresses anterior to posterior from 3 to 5 dpf (**Fig. 2E & F**), suggests that supernumerary neurons were generated sooner in the anterior forebrain, many of which no longer express PSA-NCAM, while the more recently generated neurons of the diencephalon still express PSA-NCAM.

Given the differential effects we find in different cell types (NSCs vs. tACs) forebrain regions (telencephalon vs. diencephalon) and stages (3 vs. 5 dpf), it is possible that the combination of timepoint, region, and NSPC subtype examined could yield contradictory results from different studies. This would explain some of the dissensus in the NMDA receptor-neurogenesis literature (see Introduction). Furthermore, this also explains why the habenula, as the most anterior region of the diencephalon, and the preoptic area, as a diencephalic structure that derives from the telencephalon embryologically, would progress ahead of other diencephalic regions in the production of supernumerary neurons.

### NMDA receptor dysfunction and neurogenesis in neurodevelopmental disorder pathogenesis

Numerous disease-associated variants in the genes encoding NMDA receptor subunits are associated with neurodevelopmental disorders such as autism, epilepsy, and schizophrenia [14–16,74,75]. How these genetic changes give rise to disease, however, is not well understood but typically has been attributed to changes in synaptic function. Dysregulated neurogenesis is a key facet of several neurodevelopmental disorders including autism [10]. Our work has demonstrated that perturbation of NMDA receptor signaling results in supernumerary neurons throughout the forebrain (**Fig. 2 & 3**). Notably, in autistic patients, a key phenotype is excessive prefrontal neuron numbers [10]. Hence, while changes to synaptic transmission may play a part in NDDs, the contribution of NMDA receptor dysfunction to neurodevelopmental disorder pathogenesis may also lie in its effect on the initial architectural events that build the brain, long before synapses even exist. Neuronal overpopulation may fundamentally change the way a developing brain is wired and subsequently lead to abnormal function and the phenotypes associated with neurodevelopmental disorders. This alternative perspective on pathogenesis may offer inroads to new treatment and preventative options not considered previously.

### Prospective Questions

The spatiotemporal aspect of neurogenesis offers a possible explanation for the progressive nature of the supernumerary neuron phenotype observed in *grin1^-/-^* fish. Still, it does not explain why we saw varied levels of reduced neurogenesis in most diencephalic regions at 3 dpf. Different developmental periods and regions rely more heavily on direct neurogenesis, wherein NSCs produce neuroblasts that promptly mature into postmitotic neurons without going through an amplifying stage. Other stages and regions rely more on indirect neurogenesis whereby neuroblasts function as amplifying cells that go through several rounds of division to increase the pool of progenitors and neurons before differentiating into postmitotic neurons [76,77,21]. This, paired with our findings at 3 dpf, might suggest that NMDA receptors exert an opposite effect on NSCs as compared to tACs. Future experiments will be needed to address potential roles of NMDA receptors in early neurogenic phases.

We also capitalized on the ontogenetic upregulation of KCC2 as a metric of neuronal maturation and found reduced expression in *grin1*^-/-^ fish. Notably, other groups have shown that KCC2 insufficiency may contribute to neurodevelopmental disorder pathogenesis [78–81] and that changes in KCC2 expression and activity may have effects on neurogenesis through its modulation of GABA signaling [82–84]. Hence, the relationship between NMDA receptor dysfunction, neurogenesis, and KCC2 expression might be more than correlative, but future studies are required to define this relationship.

## Conclusion

Our work contributes to a more refined understanding of the regulatory role of NMDA receptors in neurogenesis. It demonstrates that the NMDA receptor function in developmental neurogenesis cannot be generalized to all NSPC subtypes, stages of development, or signaling modes. Rather, the effect on NPCs is distinct from that on NSCs, is concentrated in the postembryonic rather than embryonic period, and is mediated by NMDA receptor ionotropic activity. Furthermore, this work suggests a relationship between NMDA receptor function and expression of the chloride transporter, KCC2, which is implicated in both neurogenesis and neurodevelopmental disorders. Hence this work highlights the intricacies of NMDA receptor function in developmental neurogenesis and offers new perspectives on how its altered function may contribute to neurodevelopmental disorders.

## Abbreviations

See table 1

## Acknowledgements

Research reported in this publication was supported by the National Institute of General Medical Sciences of the National Institutes of Health under Award Number K12GM102778 to A.J.N, the National Science Foundation Graduate Research Fellowship under grant Number 1839287 to A.J.N, a Stony Brook University Presidential Dissertation Completion Award to A.J.N., a Simons Summer Fellowship to K.M., Stony Brook University URECA summer fellowships to B.B and O.M., NIH grants to L.P.W (R01SNS088479), H.I.S and L.P.W (5R03HD101767-02). We thank Dr. James Napoli for his assistance with R applications. Special thanks to Wendy Akmentin for technical expertise and assistance, Dr. Alice Powers for critical comments on the manuscript, Dr. Bernadette Holdener and Dr. Mary Kritzer for technical advice, Dr. Shaoyu Ge for sharing reagents, and the many undergraduate students who assisted with fish care. The content of this publication is solely the responsibility of the authors and does not necessarily represent the official views of the National Institutes of Health, National Science Foundation, or Stony Brook University.

## Statements and Declarations

### Funding

This work was supported by the National Institute of General Medical Sciences of the National Institutes of Health under Award Number K12GM102778 to A.J.N, the National Science Foundation Graduate Research Fellowship under grant Number 1839287 to A.J.N, a Stony Brook University Presidential Dissertation Completion Award to A.J.N., a Simons Summer Fellowship to K.M., Stony Brook University URECA summer fellowships to B.B and O.M., NIH grants to L.P.W (R01SNS088479), H.I.S and L.P.W (5R03HD101767-02).

### Competing Interest

The authors declare no competing financial interests.

### Author Contributions

The authors confirm contribution to the paper as follows: study conception and design: Amalia J. Napoli, Howard I. Sirotkin, and Lonnie P. Wollmuth; generation of *grin1*^-/-^ mutant line: Josiah D. Zoodsma; data collection: Amalia J. Napoli, Stephanie Laderwager, Bismi Biju, Olgerta Mucollari, Sarah K. Schubel, Christieann Aprea, Aaliya Sayed, Kiele Morgan, Annelysia Napoli, Stephanie Flanagan; analysis and interpretation of results: Amalia J. Napoli, Howard I. Sirotkin, and Lonnie P. Wollmuth; figure construction: Amalia J. Napoli and Annelysia Napoli; draft manuscript preparation: Amalia J. Napoli, Howard I. Sirotkin, and Lonnie P. Wollmuth. All authors reviewed the results and approved the final version of the manuscript.

### Data Availability

Raw datasets generated and analyzed during the current study will be made publicly available in the OSF data repository as the manuscript is processed.

### Ethics Approval

All experimental work contained herein was approved and conducted in accordance with the Stony Brook University Institutional Animal Care and Use Committee (IACUC).

---

**Supplementary Figure 1.**
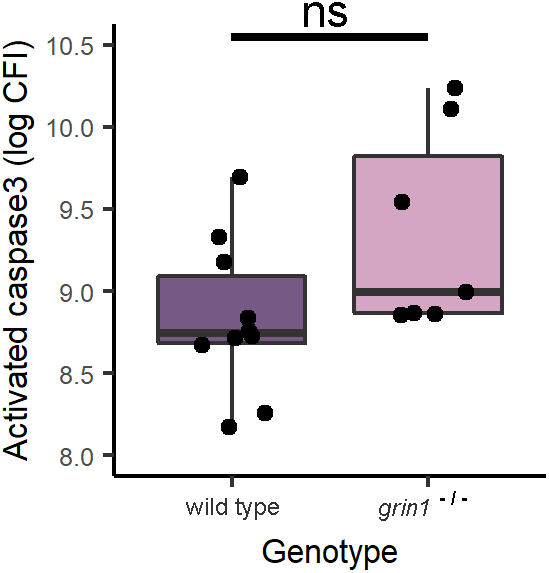
Supernumerary neurons in the forebrain of *grin1*^-/-^ fish do not result from a failure of programmed cell death. CFI results for assay of activated caspase 3 in 3 dpf fish showing no statistical difference between the level of programmed cell death in wild-type and *grin1*^-/-^ fish. Wild type n = 10, *grin1*^-/-^ n = 7.

**Supplementary Figure 2.**
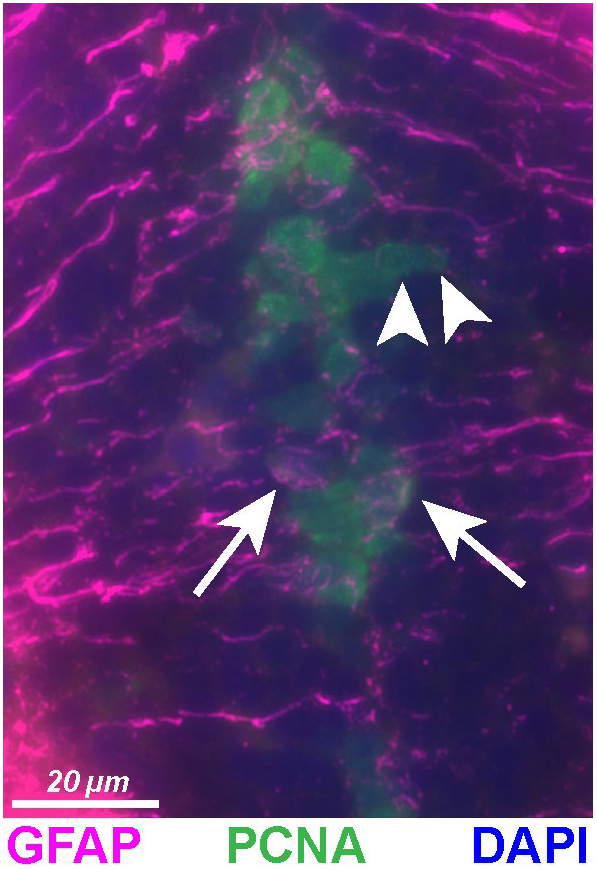
Identification of radial glia and transit amplifying cells with GFAP and PCNA Expression. Wild-type 3 dpf fish expressing GFAP and PCNA with DAPI counterstain. Example GFAP^+^ and PCNA^+^ cells at PVZ designated as RGCs expressing (arrows). Example GFAP^-^ and PCNA^+^ abventricular cells designated as tACs (arrowheads).

## Notes

### Competing Interest Statement

The authors have declared no competing interest.

### Summary of Updates

Title change. Revised introduction.

